# Spatiotemporal Patterns and Structural Substrates of Individual Functional Variability in Youth

**DOI:** 10.64898/2026.06.16.730254

**Authors:** Zekun Yang, Xiaoxi Dong, Debin Zeng, Lei Chu, Yirong He, Jichang Zhang, Qiongling Li, Yiyu Zhang, Lianglong Sun, Jing Cong, Zaixu Cui, Xiuying Wang, Shuyu Li

## Abstract

Youth is a period of emerging individuality and extensive neural remodeling, yet how functional brain individuality is organized across development remains unclear. Prior work has often conflated variability in functional topography with variability in functional connectivity, highlighting the need to examine them separately to better understand how functional individuality relates to brain structure and cognition. Here, using individualized functional parcellation in two large multimodal developmental cohorts (total N = 1211), we separately quantified variability in individualized functional parcellation (vIFP) and in functional connectivity (vFC). Both forms of variability were organized along the sensorimotor–association axis and were greatest in the association cortex. During development, vIFP increased significantly with age at both regional and whole-brain level, whereas vFC showed regionally specific age effects but no significant whole-brain increase. Both trajectories exhibited a common mid-adolescent inflection (14–16 years), marking a window of accelerated functional individualization. Despite shared spatial and temporal organization, vIFP and vFC showed dissociable associations with brain structure and cognition. vIFP was more strongly coupled to structural variability, whereas vFC was more strongly associated with cognitive variability. These findings reveal convergent and divergent developmental principles of topographic and connectional functional variability, highlighting their complementary roles in structural constraints and cognitive specialization.

## INTRODUCTION

Youth constitutes a pivotal neurodevelopmental window during which multiple cognitive, affective, and psychosocial capacities are established or refined^1–3^. Previous studies have shown that brain functional organization continues to develop throughout youth and reaches maturity in early adulthood^4–6^, and that individualized features of brain functional organization are linked to complex cognitive abilities, including executive control and social cognition^7–13^. However, adolescent maturation is highly heterogeneous, with substantial inter-individual variability in behavioral and neurobiological trajectories^14^. This phenotypic diversity likely reflects individual differences in anatomical structure (including cortical morphology and white matter microstructure) and functional networks^14–21^. Thus, characterizing developmental changes in individual functional variability could provide quantifiable evidence for the individualized maturation of the human brain and further enhance our understanding of the diversity in cognition and behavior during youth. These insights may facilitate the clinical translation of neuroscience findings toward precision medicine, tailored education, and personalized neuromodulation^13,14,16,22–24^.

Functional brain development involves the progressive refinement of cortical parcellation, whereby functional interactions become stronger within the same cortical region and weaker between distinct regions, leading to increasingly differentiated functional boundaries. Such refinement indicates that developmental variation in functional organization can be characterized along two complementary dimensions^18,25^: functional connectivity (FC) architecture, represented by FC profiles^13,17,26–31^, and functional topography, which captures the spatial layout of functional parcels^7,10,15,28,32,33^. From birth onward, both forms of functional variability are heterogeneously distributed across the cortex along the sensorimotor–association (S-A) axis of neurodevelopment^7,10,27–29,32,34,35^. More recent work extended these region-wise findings to edge-wise FC variability, identifying a stable connectional axis and characterizing developmental changes in edge-wise variability and the axis, whose slope flattened during youth^30^. Developmental changes in functional variability during youth are intrinsically linked to phenotypic diversity, explaining the variance in executive control during early adolescence and the emergence of affective and psychotic symptoms in later stages^13,30,36^. Nevertheless, existing developmental studies have only partially characterized these two forms of individual functional variability. In particular, continued reliance on population-level atlases limits the detection of subject-specific functional features^7,10,13,28,30,37,38^, while the developmental trajectories of variability in functional topography and FC profiles remain insufficiently understood^7^. To address this gap, we used an individualized functional parcellation approach to delineate the spatiotemporal patterns of inter-individual variability in functional topography and connectivity across youth.

Brain structural architecture serves as the physical substrate for functional dynamics. Youth represents a period of profound neurobiological flux, characterized by ongoing cortical remodeling, myelination, and white-matter maturation^34,39–41^. These developmental processes may shape individual variability in functional organization through multiple structural dimensions, including cortical morphology, white-matter connectivity, and myelin content^42–46^. In adults, structural variability has been shown to capture aspects of functional organization^45,47^, partly through regional structure-function coupling^48,49^. Whole-brain comparisons of spatial similarity between structural and functional variability have revealed similar cortical patterns^8,30,49,50^. Crucially, most prior developmental studies relied on population-averaged atlases to examine structure-function relationships^8,30,49^. While such approaches successfully delineate how functional profiles vary at fixed anatomical coordinates across individuals, they inherently fail to address the inverse question regarding how structural substrates differ within true homologous functional boundaries. Therefore, examining the relationship between brain structure and function within subject-specific functional boundaries is necessary to clarify the structural substrates of individual functional variability during development.

In this study, we systematically investigated the spatiotemporal patterns, structural substrates, and cognitive relevance of inter-individual functional variability. We used multimodal neuroimaging data from the Human Connectome Project Development (HCP-D)^51,52^ as the discovery cohort and tested the generalizability of the resulting functional findings in an independent replication sample from the Philadelphia Neurodevelopmental Cohort (PNC)^53^. Using an individualized functional parcellation (IFP) framework, we quantified inter-individual variability in functional parcellation (vIFP) and functional connectivity (vFC) and characterized their cortical distributions and developmental trajectories. To identify the structural substrates of functional variability, we quantified variability in white-matter structural connectivity, morphometric similarity networks, and cortical myelin distribution within individual-specific functional boundaries. We then assessed global spatial correspondence and regional coupling between structural and functional variability across homologous functional regions. Finally, we examined the cognitive relevance of vIFP and vFC by testing their ability to predict inter-individual variability in crystallized, fluid, and total intelligence.

## RESULTS

We analyzed multimodal neuroimaging data from 601 participants (aged 6–22 years) in the HCP-D discovery cohort^51^. Using an iterative optimization framework that identifies homologous functional regions across subjects^28,54^, we generated individualized functional parcellations (IFPs) comprising 87 discrete regions per subject. We quantified topographic variability (vIFP) as one minus the Dice overlap between homologous parcels across individuals. To maintain functional homology for multimodal indicators, we utilized the same subject-specific functional boundaries to extract a comprehensive suite of structural and functional features, including functional connectivity (FC), white-matter structural connectivity (SC), morphometric similarity networks (MSN), and myelin distribution (MD). We then quantified their respective inter-individual variabilities (vFC, vSC, vMSN, and vMD). To map the maturational trajectories of these features, a cross-subject sliding-window approach and generalized additive models (GAMs) were then used to characterize nonlinear age-related trajectories in functional and structural variability throughout youth. At the regional level, we quantified coupling between functional and structural variability within homologous regions to investigate the structural substrates of functional variability. We also used multivariate ridge regression models to predict pairwise differences in crystallized, fluid, and total intelligence from vIFP and vFC, both separately and jointly, thereby assessing their relative cognitive relevance (Fig. 1). Finally, we tested whether the principal spatial and developmental patterns of vIFP and vFC could be replicated in an independent cohort of 610 participants aged 8–23 years from the Philadelphia Neurodevelopmental Cohort (PNC).^53^.

**Figure 1.**
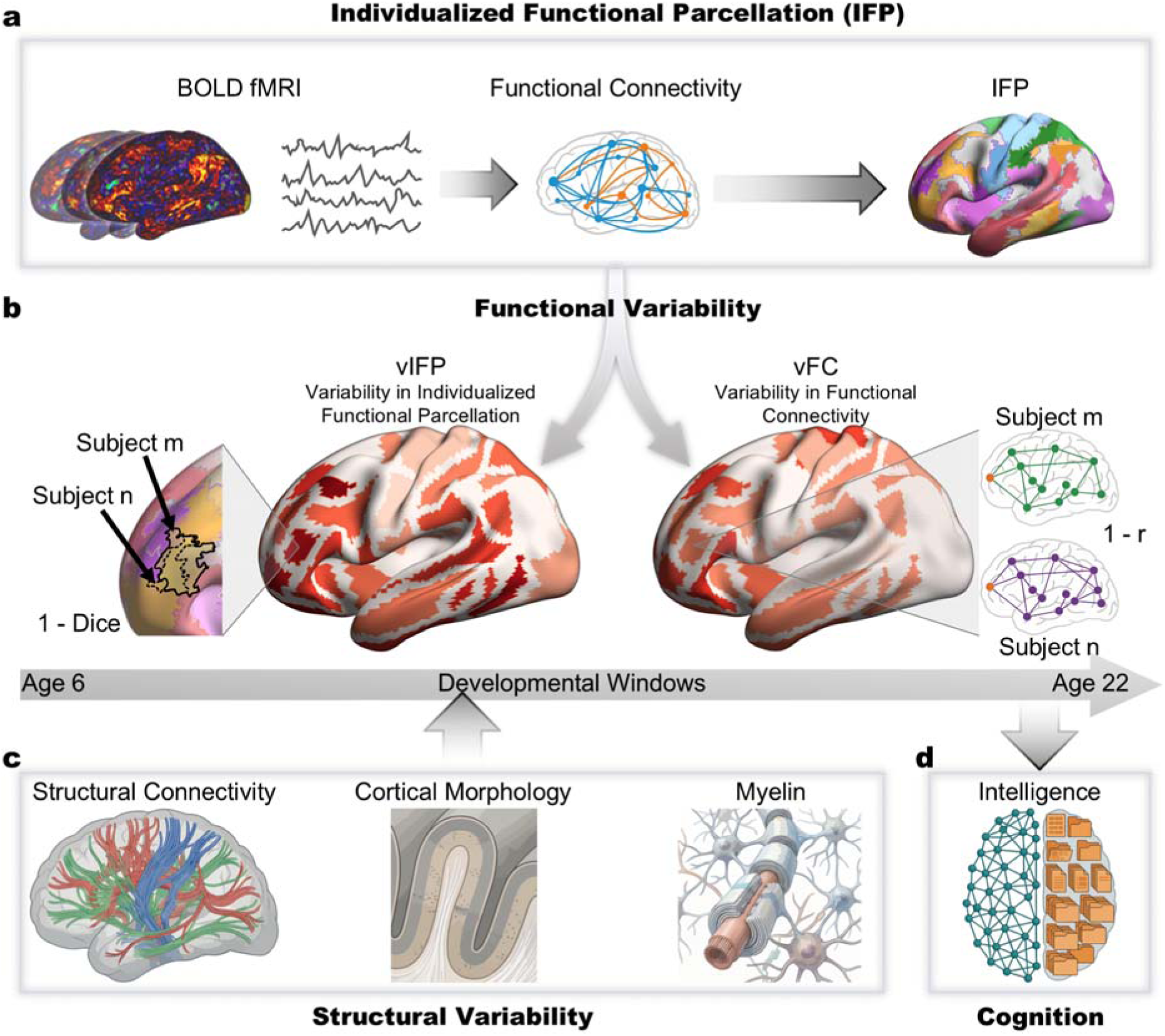
A multi-scale framework for quantifying the spatiotemporal patterns and structural substrates of functional variability. **a. Construction of individualized functional parcellation (IFP).** IFPs were derived from resting-state functional MRI (rs-fMRI) data for each participant to capture subject-specific functional topographies. **b. Dissociating components of functional variability.** Inter-individual functional variability was separated into two distinct dimensions: variability in functional parcellation (vIFP) and variability in functional connectivity (vFC). vIFP was quantified by calculating the spatial overlap (Dice coefficient) of parcellations across individuals, while vFC was measured by comparing the similarity of IFP-based connectivity profiles. The developmental trajectories of these components were modeled across a cohort spanning 6 to 22 years. **c. Multiple structural substrates.** To identify the neurobiological substrates of functional individuality, variability in white-matter structural connectivity (vSC), morphometric similarity networks (vMSN), and myelin distribution (vMD) was quantified within IFP-defined boundaries. Spatial and developmental coupling between structural and functional variability was assessed to determine the structural underpinnings of brain individuality. **d. Cognitive significance.** The association between individual functional variability and higher-order cognition (e.g., intelligence) was examined, establishing the link between brain individuality and cognitive phenotypes during youth.

### Spatiotemporal Patterns of Individual Functional Variability

The IFP framework captured inter-individual variations in the location, shape and surface area of cortical functional organization. On the inflated cortical surface (Fig. 2a), the face motor regions of three representative subjects were localized within a consistent cortical vicinity but exhibited pronounced differences in boundary geometry and surface area. This topographic heterogeneity persisted across all developmental stages and remained evident even between age-matched individuals. Furthermore, on the pial surface (Fig. S1), these face motor regions displayed significant morphological divergence across subjects, primarily driven by the substantial inter-individual variability in cortical folding patterns.

**Figure 2.**
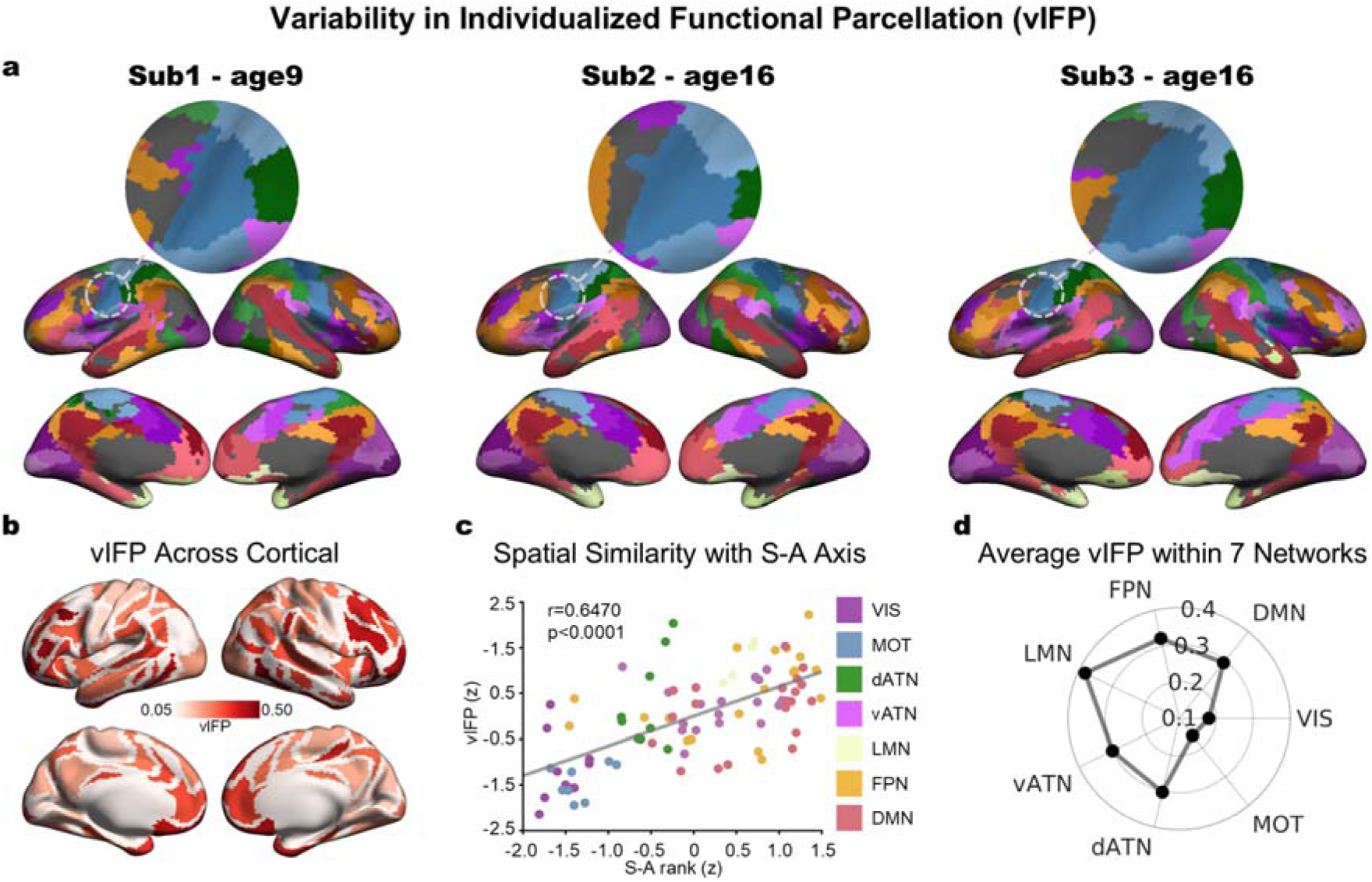
Variability in individualized functional parcellation (vIFP). **a.** IFPs from three participants of different ages. Circles highlight the facial motor area. **b.** Cortical distribution of vIFP across 601 HCP-D participants. **c.** Correlation between regional vIFP and rank along the S-A axis, with point colors indicating functional networks. **d.** The spider diagram shows the average vIFP within Yeo’s 7 networks. VIS, visual network; MOT, motor network; dATN, dorsal attention network; vATN, ventral attention network; LMN, limbic network; FPN, frontoparietal network; DMN, default mode network.

The homologous functional regions identified here covered nearly the entire cerebral cortex (Fig. S2a). To evaluate the reliability of the individualized functional parcellation approach, we performed a split-half analysis by independently dividing each participant’s BOLD fMRI data into two time-series segments. The resulting IFPs demonstrated high intra-subject reproducibility (mean Dice coefficient = 0.8176; Fig. S2b), with nearly all regions exhibiting Dice values exceeding 0.7.

We delineated the regional landscape of inter-individual topographic diversity by constructing a cortical map of vIFP (Fig. 2b). vIFP was heterogeneously distributed across the cortex, exhibiting a clear hierarchical progression aligned with the brain’s functional organization. Specifically, vIFP was markedly heightened within transmodal association cortices and minimal within primary sensorimotor regions. System-level analysis further corroborated this pattern, with the sensorimotor system demonstrating the lowest vIFP, followed by the visual system, whereas the default mode, attentional, and frontoparietal control networks exhibited substantially greater variability (Fig. 2d). The limbic network emerged as the most variable system, displaying the highest vIFP. This spatial heterogeneity may reflect the fundamental hierarchical organization of the human brain, as evidenced by a robust correlation between the cortical distribution of vIFP and the S-A axis (r = 0.6470, p < 0.0001; Fig. 2c). Furthermore, the observed spatial and system-level distribution of vIFP was highly consistent with prior evidence using non-negative matrix factorization (NMF) ^7^ to quantify individual functional topography in youth, reinforcing the robustness of our IFP-derived metrics in capturing individualized functional organization.

Our analysis revealed that vIFP underwent a significant age-related increase during youth. To investigate the developmental changes in individual variability, we calculated vIFP for each group and implemented a paired cohort comparison between children (6–12 years, n=147) and young adults (>18 years, n=147). We observed a striking conservation of the spatial landscape of vIFP across development (r = 0.9693, p < 0.001; Fig. S3). The vIFP values in the young-adult group were significantly higher than those in the child group (paired t-test, p=0.0268). Our results suggested that the hierarchical organization of individual differences along the S-A axis was established early and maintained into maturity.

To delineate the temporal evolution of individualized brain organization, we implemented a high-resolution, cross-subject continuous sliding window approach (window size = 20, step size = 10; resulting in 59 overlapping age groups). We modeled the developmental trajectories of both regional and global vIFP using GAMs to capture non-linear maturation while controlling for sex and in-scanner head motion. Our analysis revealed a significant age-related increase in global mean vIFP during youth (delta R^2^ ANOVA p < 0.0001; Fig. 3a). A total of 54 regions exhibited significant age-related effects, with the majority showing a progressive expansion of vIFP (Fig. 3c, d). Despite the magnitude shifts in individualization, the spatial distribution of vIFP remained anchored to the S-A axis throughout the entire age range (Fig. 3c). The most pronounced developmental effects were localized within the dorsal and ventral attention networks (dATN/vATN) (Fig. 3c, d). These networks occupy a pivotal transitional position within the cortical hierarchy, bridging unimodal sensory systems and transmodal association networks. To pinpoint critical maturational windows, we derived the growth rate of vIFP as the first derivative of its age-related trajectory. This revealed a prominent maturational turning point at approximately age 14 (Fig. 3g). Before this mid-adolescent transition, vIFP exhibited small growth rates in most regions and regionally specific patterns, with modest declines observed in regions such as the supplementary motor area, Broca’s area, and the anterior cingulate cortex, which are often associated with early refinement of language and executive control^55^. In contrast, the period after age 14 was characterized by an accelerated increase in topographic variability with larger growth rates in nearly all regions.

**Figure 3.**
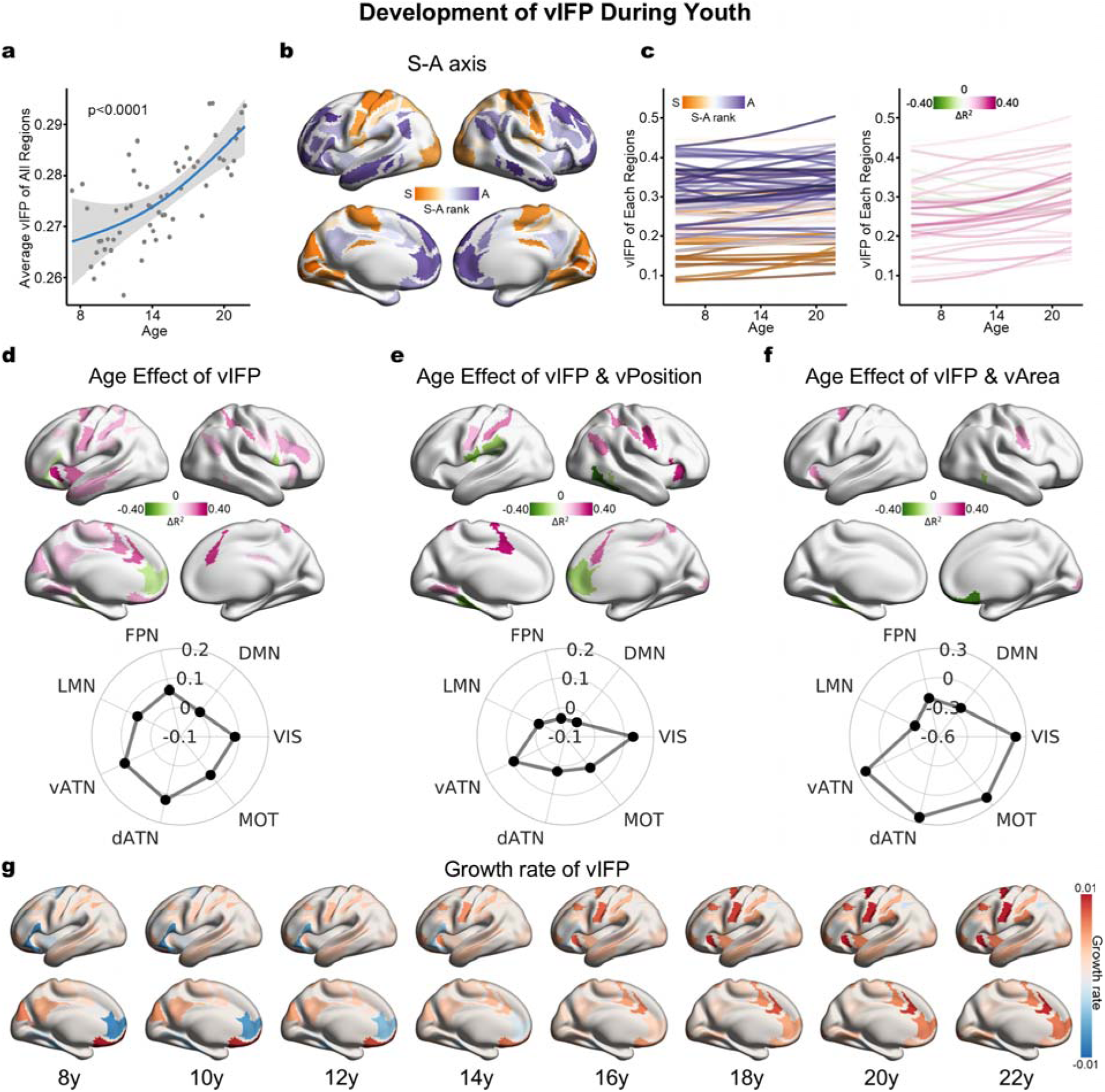
Variability in individualized functional parcellation (vIFP) among youth. **a.** Developmental trajectory of average vIFP across the whole brain. **b.** The sensorimotor-association (S-A) axis in neurodevelopment^34^. **c.** The trajectory curves of vIFP with age; each line represents a region. The x-axis denotes age, and the y-axis denotes vIFP. The color of the curves in the left figure indicates the position of the ROI on the S-A axis. The color of the curves in the right figure reflects the age effects (delta R²) on vIFP; pink indicates an increase with age, green indicates a decrease. **d.** Regions with significant age effects on vIFP, where color represents the magnitude and direction of the age effect. **e.** Regions showing significant age effects on vIFP and significant age effects on position variability (vPosition). **f.** Regions with significant age effects on both vIFP and areal variability (vArea). Average age effects within Yeo’s 7 functional networks are depicted in the radar chart below. **g.** The growth rate of vIFP was calculated as the first derivative (with respect to age) of the developmental trajectory fitted by a generalized additive model (GAM).

To elucidate the factors that may drive developmental changes in vIFP, we characterized vIFP using two measures: positional variability (vPosition), which reflects the spatial displacement of homologous functional regions, and areal variability (vArea), which reflects inter-individual differences in their surface area. vPosition was quantified as the mean cross-subject Euclidean distance between homologous parcel centroids, while vArea captured the variance in region size across individuals. Our analysis revealed a striking dissociation in the maturational trajectories of these two measures. Among the 54 cortical regions exhibiting significant age-related effects in vIFP, the majority (34 regions; 63.0%) showed significant age effects in vPosition (Fig. 3e). In contrast, age-related changes in vArea were sparse, with only 7 regions (13.0%) reaching significance (Fig. 3f). These results provided evidence that the individualized maturation of functional organization during youth was predominantly driven by positional changes.

To derive an accurate representation of functional architecture, we calculated individualized functional connectivity (FC) profiles within the subject-specific boundaries defined by our individualized functional parcellation (IFP). We quantified inter-individual variability in these FC profiles (vFC; Fig. 4a) and assessed age effects using the same methodology described previously (Fig. 4b, d). To control for the impact of noise and other technical confounds on inter-subject vFC estimates, inter-subject vFC was corrected by regressing out mean intra-subject variability using a strategy similar to that described by Mueller and colleagues^29^. Only participants with four resting-state fMRI runs were included, yielding a final sample of 579 individuals. Compared to conventional registration methods based on population-level atlases, the IFP framework reduced spurious inter-subject variability in functional connectivity architecture (Fig. S4a). By explicitly accounting for subject-specific functional boundaries, our IFP method ensured that FC profiles reflected functional synchronization rather than registration artifacts.

**Figure 4.**
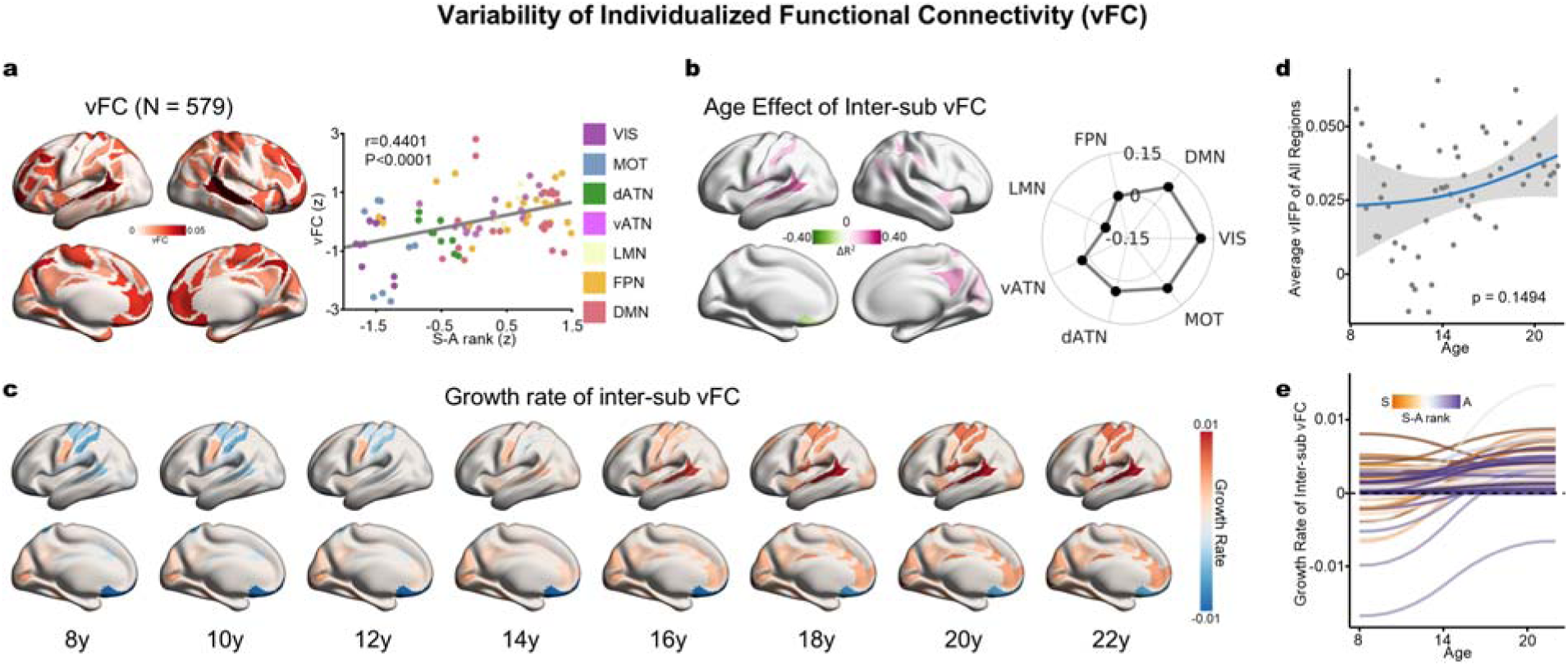
Inter-individual variability in individualized functional connectivity (vFC). **a.** Distribution of inter-individual vFC across the cortex, and its spatial similarity to the S-A axis. **b.** Regions with significant age effects on inter-subject vFC (delta R²). **c.** Regional growth rates of inter-subject vFC, calculated as the first derivative of growth trajectories fitted using GAMs. **d.** Developmental trajectory curves of average inter-subject vFC across the whole brain, with scatter points representing sliding windows. The age effect was not significant (delta R² ANOVA p = 0.1494). **e.** Growth-rate curve for inter-subject vFC (first derivative of developmental trajectories).

vFC was non-uniformly distributed across the cortex, and variability was highest within the association cortices and significantly attenuated within the sensorimotor cortices, aligning with the S-A axis of neurodevelopment (r = 0.4401, p < 0.0001; Fig. 4a). We identified 39 regions exhibiting significant age-related changes in vFC (delta R² ANOVA FDR-corrected p < 0.05; Fig. 4b). The most pronounced developmental changes were localized within the visual network (VIS) (average absolute delta R^2^=0.1053) and the default mode network (DMN) (average absolute delta R² = 0.0778), followed by the motor network (MOT) (average absolute delta R² = 0.0730), the dorsal attention network (dATN) (average absolute delta R² = 0.0398), and the ventral attention network (vATN) (average absolute delta R² = 0.0270, Fig. S5). However, the age effect on global mean vFC across the entire cortex did not reach significance (delta R² ANOVA p = 0.1494).

The growth rate of inter-subject vFC followed a non-linear trajectory that closely mirrored the spatiotemporal evolution of vIFP, characterized by a maturational inflection point at approximately age 14 (Fig. 4c). In contrast to this inter-individual expansion, intra-subject vFC exhibited a sustained and monotonic decline throughout youth (Fig. S4b, c). The divergent developmental trajectories between inter- and intra-subject variability underscored a profound neurodevelopmental change. As the nervous system matured, it concurrently shaped a more robust foundation and a highly individualized functional profile.

### Dissociable Developmental Trajectories of Structural Variability

To uncover the structural substrates underlying personalized functional organization, we utilized the subject-specific boundaries defined by our IFP to extract a tripartite set of structural features. First, we quantified inter-individual variability in white-matter structural connectivity (vSC), derived from high-resolution probabilistic tractography to capture the complex fiber architecture of the human connectome. We then characterized inter-individual variability in morphometric similarity networks (vMSN), representing patterns of inter-regional similarity in cortical morphometric profiles. Finally, we assessed the variability in cortical myelin distribution (vMD) by computing the Kruskal-Wallis H statistic on T1/T2 ratios across individuals. Details are provided in the Methods section.

Spatially, the inter-individual variability of these structural metrics exhibited distinct distribution patterns across the entire cortex. Both vMSN and vSC displayed a clear hierarchical pattern, with heightened variability within the association cortices and relative conservation in primary regions (Fig. 5a). Specifically, vMSN exhibited a robust positive alignment with the S-A axis (r=0.5770, p<0.0001; Fig. 5a). While the correlation between vSC and this hierarchical axis was more moderate, it remained statistically significant (r=0.2710, p=0.0111; Fig. 5a). At the system level, both structural features peaked within the frontoparietal control network (FPN) and reached their minima in the visual (VIS) system (Fig. S6a, b). In a departure from cortical morphology and structural connectivity, the spatial distribution of vMD showed no significant alignment with the S-A axis (r=0.1277, p=0.2575; Fig. 5a). Instead, vMD reached its maximum in the dorsal and ventral attention (dATN/vATN) and limbic (LMN) networks, while remaining the lowest within the MOT and VIS systems (Fig. S6c). This observed spatial dissociation challenged previous models derived from population-based templates, which consistently reported strong hierarchies for vSC, vMSN, and vMD^49^. Our findings, derived from the IFP framework, offered a novel perspective that refined our understanding of structural variability.

**Figure 5.**
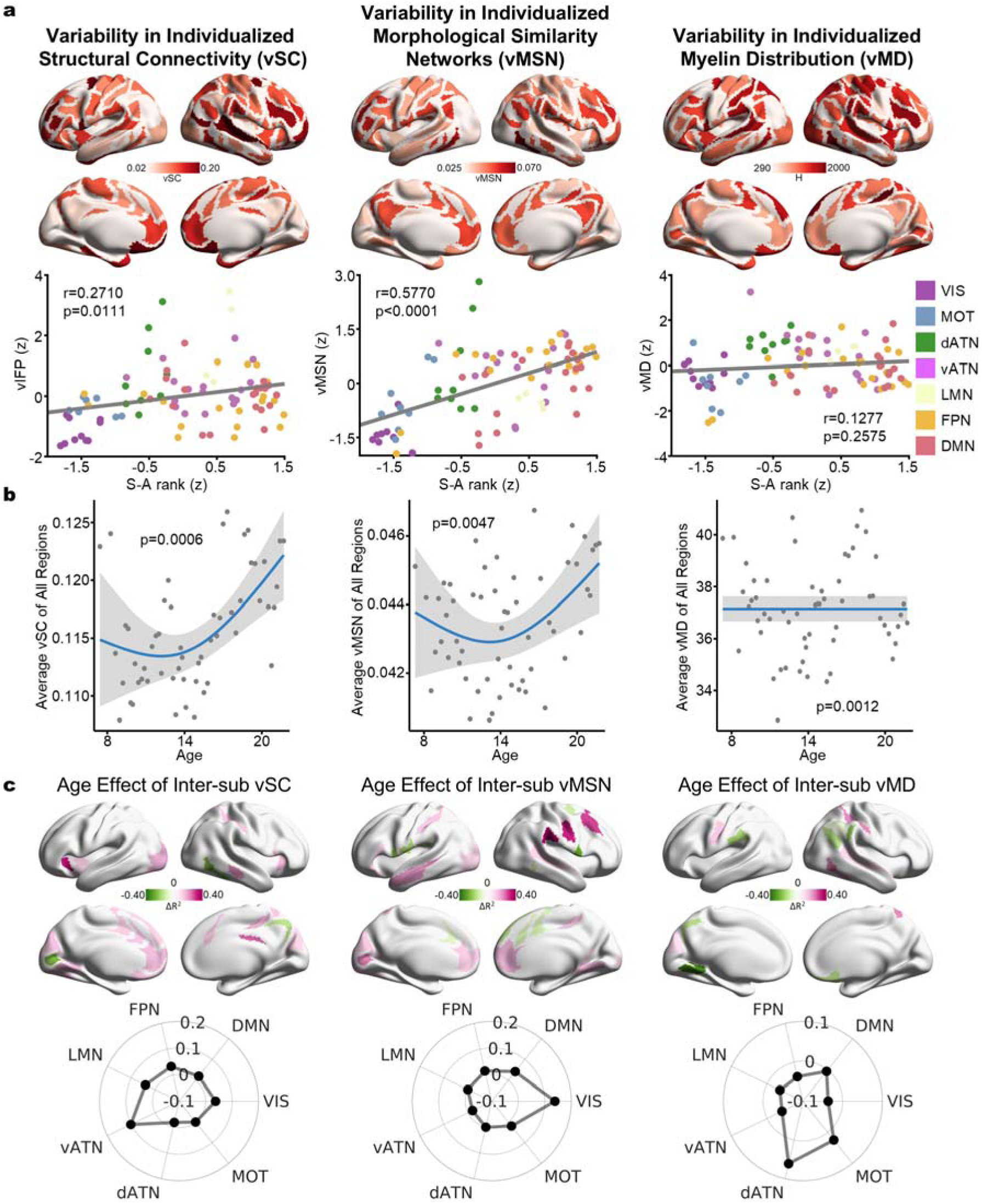
Inter-individual variability of structural features. **a.** Inter-individual variability for three IFP-based structural features: variability in white-matter structural connectivity (vSC), variability in morphometric similarity networks (vMSN), and variability in myelin distribution (vMD). **b.** Developmental trajectories of average structural variability across the whole brain. The p-value from the GAM fit is indicated. **c.** Regions showing significant age effects in structural variability. Colors indicate the magnitude (delta R^2^) and direction of developmental effects.

Beyond spatial patterns, the three structural metrics exhibited highly divergent spatiotemporal trajectories throughout youth. Averaged across the whole brain, both vSC (delta R^2^ ANOVA p = 0.0006) and vMSN (delta R² ANOVA p = 0.0047) demonstrated significant age-related increases, following a non-linear, U-shaped trend. Both trajectories were characterized by an initial slow decline and a subsequent accelerated rise after the inflection point at approximately age 14 (Fig. 5b). In contrast, vMD showed no significant whole-brain age effect. Although adding age improved model fit (delta R² ANOVA p = 0.0012), the smooth term was not significant (p = 0.6261), arguing against a robust developmental trajectory (Fig. 5b). The regional drivers of these trajectories further underscored this structural dissociation. The age-related growth of vSC was primarily anchored within the vATN and LMN (Fig. 5c). Conversely, the expansion of vMSN was more globally manifested within the VIS and DMN systems. For vMD, with the exception of a positive effect in the dATN and MOT, age effects across the remaining six systems were entirely negative (Fig. 5c). These system-level trends were robustly recapitulated at a fine-grained level. We identified a substantial number of regions exhibiting significant age-related shifts: 53 regions for vSC, 60 regions for vMSN, and 51 regions for vMD (delta R² ANOVA FDR-corrected p<0.05; Fig. 5c). Crucially, these ROI-level variations showed high spatial alignment with the whole-brain trends, with vSC and vMSN showing nearly universal increases across significant regions. We assessed the spatial similarity between the age-effect (delta R²) maps of functional and structural variability and found a significant correlation only between the vIFP and vSC maps (r = 0.3902, p = 0.0002; Fig. S7).

To further elucidate the temporal dissociation of structural variability, we derived annualized growth rates by calculating the first derivatives of the GAM-fitted developmental trajectories for each structural phenotype. Paralleling our functional findings, structural variability exhibited a pivotal maturational window between ages 14 and 16 years. The specific growth trends diverged markedly across metrics (Fig. S8). The growth curves of vSC and vMSN captured a distinct period of accelerated individual differentiation. Within the 14–16-year window, the growth rates in numerous regions underwent a sign reversal, shifting from negative to positive. Following this inflection, both vSC and vMSN entered a phase of accelerated expansion that was most prominent within the association cortex (Fig. S8a, b). In contrast, the growth rates of the microstructural phenotype vMD exhibited substantial regional heterogeneity throughout youth, with most values hovering near zero (Fig. S8c). This indicated that myelin distribution remained relatively stable within homologous functional boundaries, while the architectures of white matter connectivity and morphometric similarity underwent a progressive divergence.

### Population-wide Coupling of Structural and Functional Individual Variability

To integrate these multidimensional facets of individual variability, we utilized the ROI-level IFP as a high-precision common reference frame. Within this framework, we systematically examined the regional coupling between the two forms of functional variability (vIFP and vFC) and variability across three structural metrics (vSC, vMSN, and vMD) (Fig. 6). We posited that greater inter-individual variability in the spatial locus and shape of IFP-defined regions would correlate with greater inter-individual variability in their underlying structural characteristics. Our experimental results provided partial support for this hypothesis, yet uncovered a complex landscape of dissociable structure-function relationships.

**Figure 6.**
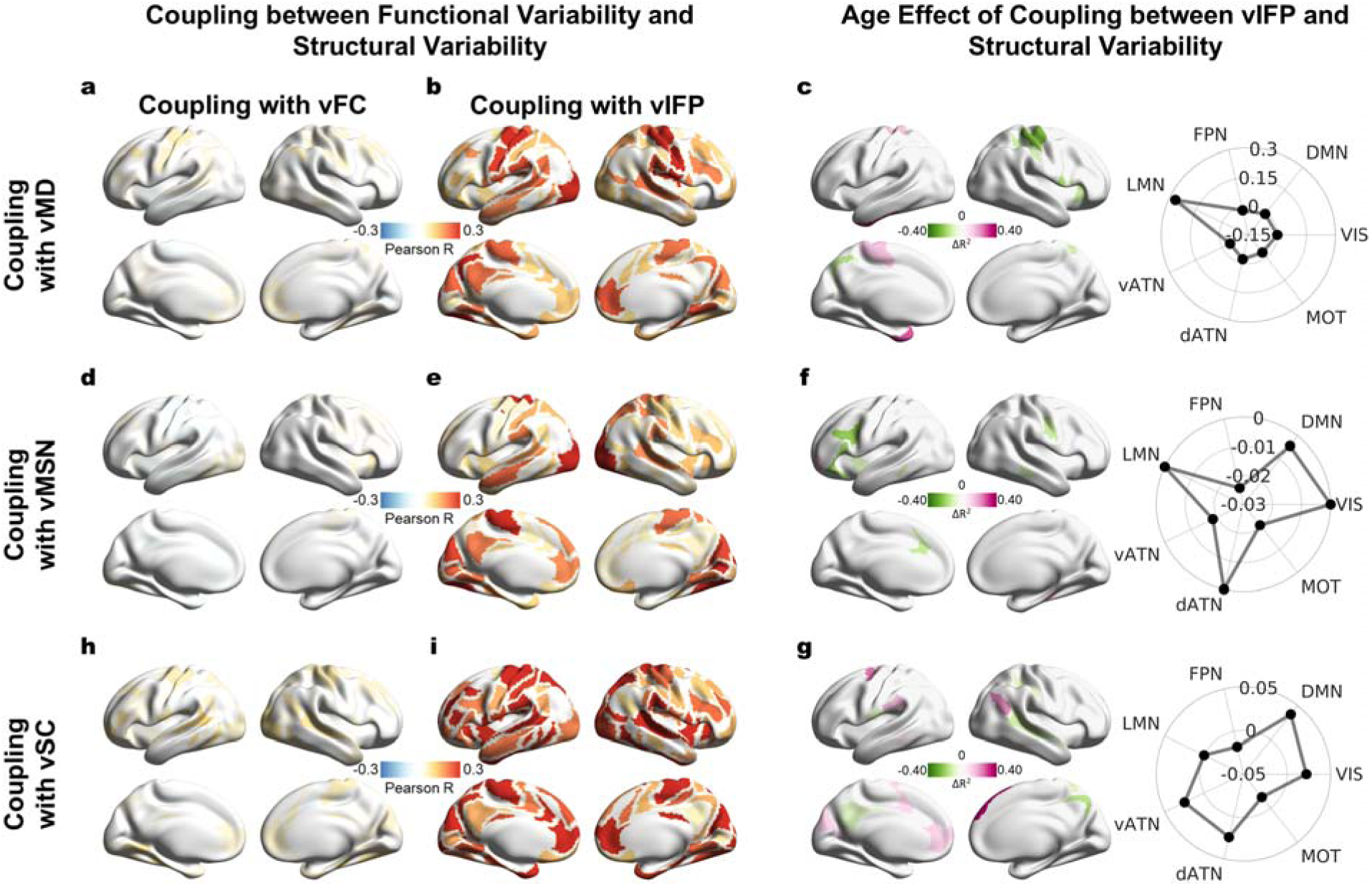
Population-wide regional coupling of structural and functional individual variability. a, b, d, e, h, **i.** Pairwise coupling of functional variabilities (vIFP and vFC) with structural variabilities (vMD, vMSN, and vSC). **c, f, g.** Age-related effects on coupling. Left panels: Regions showing significant age effects in the coupling between vIFP and structural variability. Right panels: Average age effects (delta R2) within 7 functional networks

Our analyses revealed a striking divergence in how structural variability supports functional variability. Specifically, we observed that structural variation served as a constraint on functional topography (vIFP) rather than functional connectivity (vFC) architecture. Across all three structural metrics, structural variability exhibited regional coupling with individual variability in functional parcellation (vIFP) (Fig. 6b, e, i). In contrast, the correlations between these structural variabilities and the variability of functional connectivity profiles (vFC) remained consistently weak (Fig. 6a, d, h), a discrepancy in coupling strength that was statistically significant (Fig. S8a). This indicated that the physical scaffolds of the brain (myelin, cortical morphology, and white matter connectivity) primarily dictated the spatial layout of functional boundaries, rather than inter-regional communication. The regional distribution of these couplings further highlighted phenotype-specific patterns. Among all structural features, the relationship between vSC and vIFP showed the strongest population-wide correlation (Fig. 6i and S8a), maintaining significant coupling even within the association cortex. Regionally, the strongest coupling between vMD and vIFP was heavily concentrated within sensorimotor areas (Fig. 6b), while the coupling between vMSN and vIFP peaked within the visual and attentional networks (Fig. 6e).

The coupling strength between structural and functional variability was not uniform across the cortex, adhering instead to a spatial hierarchy. For all three structural metrics, the structure–function coupling exhibited a significant negative correlation with the S-A axis (Fig. S9b–d). This hierarchical gradient was most pronounced for morphometric similarity (vMSN: r=−0.5125, p=3.89×10^-7^, Fig. S9c) and myelin distribution (vMD: r=−0.4980, p=9.18×10^-7^, Fig. S9b), while remaining significant for white-matter connectivity (vSC: r=−0.2126, p=0.0480, Fig. S9d). This uniform negative gradient provided evidence of a spatially patterned decoupling effect along the cortical hierarchy. In primary cortices, functional topography was constrained by the underlying cortical morphology and myelination, reflecting the developmental programming and lower plasticity of these systems. Conversely, higher-order association cortices demonstrated a clear release from these structural tethers. Within these association cortices, functional topography exhibited extensive inter-individual divergence even when the underlying structural variability remained restricted.

The coupling between structural and functional variability underwent profound remodeling during youth. The developmental effects (delta R^2^) predominantly reflected declines in coupling strength and exhibited distinct network-specific patterns across different structural metrics. The vMSN-vIFP coupling exhibited the most uniform maturation, with 55 regions showing significant age-related changes (delta R^2^ ANOVA FDR-corrected p<0.05). Across all seven functional networks, the mean age effect remained consistently negative (Fig. 6f), indicating a progressive decoupling of cortical morphology from functional topography. In contrast, both vMD-vIFP and vSC-vIFP couplings displayed more complex developmental trajectories. Significant developmental changes in vMD–vIFP coupling were identified in 43 regions (delta R² ANOVA FDR-corrected p<0.05). Although most networks followed the trend toward decoupling, a distinct positive age effect emerged within the LMN (Fig. 6c). For vSC–vIFP coupling, 56 regions showed significant age effects (delta R² ANOVA FDR-corrected p<0.05), with positive mean age effects within the attention networks, DMN, and VIS (Fig. 6g). These findings indicated that structure-function relationships underwent a complex, multimodal recalibration during youth, trending largely toward decoupling. When coupling strength was averaged across the whole brain, no significant age effects were observed between vIFP and any of the three structural metrics (Fig. S9e–g). By ensuring fine-grained functional homology across participants, our framework effectively unmasked spatially dissociable developmental principles that may be obscured by global, whole-brain averages or population-level templates.

### Functional Variability Predicts Cognitive Variability

To investigate the behavioral relevance of functional individuality, we analyzed crystallized, fluid, and total intelligence scores from the NIH Toolbox within the HCP-D dataset. Cognitive variability was rigorously quantified as the absolute pairwise difference in test scores between participants. We then implemented a multivariate ridge regression framework to predict this cognitive variability from regional functional variability profiles. Given that functional topography and connectivity architecture have been shown to capture dissociable neurobiological information, we first evaluated the independent predictive utility of vIFP and vFC. To further examine whether these two dimensions provided synergistic information, we developed a joint model that harnessed both vIFP and vFC. The performance of all predictive models was validated through 100 iterations of randomized two-fold cross-validation.

Our multivariate predictive modeling revealed that functional variability predicted cognitive variation, as evidenced by a significant correlation between predicted and observed cognitive variability across all domains (permutation testing, 100,000 iterations, p<0.0001). However, we observed a striking divergence in the predictive utility of the two functional dimensions. While vIFP successfully encoded cognitive information, its predictive accuracy for crystallized, fluid, and total intelligence was significantly lower than that of vFC (paired t-test, p<0.0001; Fig. 7a). When predicting fluid and total intelligence, there were no significant differences in prediction accuracy between using vFC alone and using both vFC and vIFP together (paired t-test, p>0.05; Fig. 7a, middle and right panels). For crystallized intelligence, the model utilizing vFC alone significantly outperformed the joint model (paired t-test, p=0.0002; Fig. 7a, left panel). The inclusion of vIFP features did not significantly improve prediction performance, suggesting a stronger association of vFC with cognitive variability.

**Figure 7.**
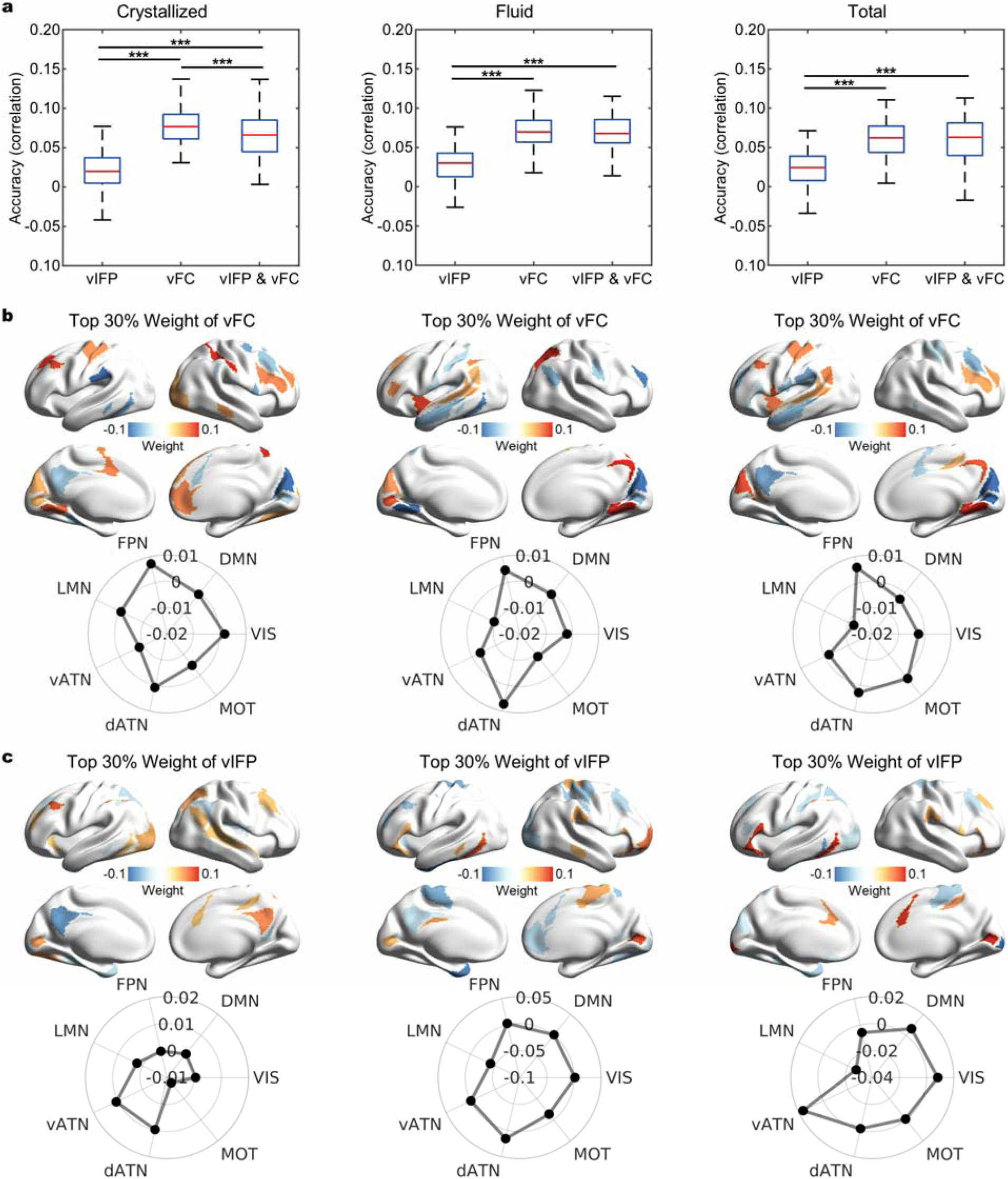
Predicting Intelligence Variability Using Functional Variability. **a.** Ridge regression was used to predict individual variability in crystallized intelligence, fluid intelligence, and total intelligence through three approaches: using vIFP alone, using vFC alone, and using both vIFP and vFC together. **b.** The top 30% of brain regions with the highest absolute weights in the ridge regression model when predicting intelligence variability using vFC alone. **c.** The top 30% of brain regions with the highest absolute weights in the ridge regression model when predicting intelligence variability using vIFP alone.

The weight distribution in the ridge regression model reflected the contributions of different brain regions to predicting intelligence. Notably, the cortical areas contributing most to cognitive prediction were non-uniformly distributed, exhibiting distinct spatial patterns for vIFP and vFC. For intelligence prediction, vIFP model weights were concentrated primarily in the vATN and dATN, whereas vFC model weights were concentrated primarily in the FPN (Fig. 7b, c). Across both functional dimensions and all intelligence domains, the most informative regions for predicting cognitive performance were predominantly anchored within the association cortex.

### Sensitivity Analysis

The 20-subject age-ordered sliding window with a 10-subject step was treated as the primary analysis. It identified regional age effects in 54 vIFP regions, 39 vFC regions, 53 vSC regions, 60 vMSN regions and 51 vMD regions after FDR correction. At the whole-brain model-comparison level, adding age improved fit for vIFP, vSC, vMSN and vMD. The whole-brain vFC effect did not reach significance.

As a sensitivity analysis, we repeated the analyses using a 10-subject age-ordered sliding window with a 5-subject step. For functional variability, the sensitivity analysis reproduced 43 of 54 vIFP regions (79.6%) and 25 of 39 vFC regions (64.1%). For structural variability, it reproduced 44 of 53 vSC regions (83.0%), 46 of 60 vMSN regions (76.7%) and 37 of 51 vMD regions (72.5%). The denser window identified 51, 57, 59, 56 and 54 significant regions for vIFP, vFC, vSC, vMSN and vMD, respectively. At the whole-brain model-comparison level, adding age improved fit for vIFP, vSC and vMD under the denser window (p = 7.20×10^-8^, 0.0020 and 0.0035). In contrast, vFC became significant (p = 0.0060), whereas vMSN retained a positive direction but did not reach whole-brain significance (p = 0.1352). These findings supported the robustness of the primary spatiotemporal patterns and identified global vFC and global vMSN as comparatively window-sensitive endpoints.

### Replication in an Independent Youth Dataset

We replicated the spatiotemporal patterns of vIFP and vFC in an independent PNC cohort of 610 participants^53^. The replication analysis was restricted to the same 87 homologous functional regions identified in HCP-D (Fig. S11). The cortical distribution of vIFP reproduced the sensorimotor-association (S-A) hierarchy observed in HCP-D, with greater variability in association cortices and lower variability in sensorimotor cortices (r = 0.5409, p < 0.0001; Fig. S11a). The whole-brain mean vIFP increased significantly with age (delta R^2^ ANOVA p < 0.0001; Fig. S11c). Of the 54 HCP-D regions with significant age effects, 33 (61.1%) were replicated in PNC under the same significance criterion (ANOVA FDR-corrected p < 0.05; Fig. S11b). Regional vIFP growth rates generally increased across development, supporting accelerated topographic individualization in youth (Fig. S11d). vFC showed a similarly reproducible spatial hierarchy, with higher variability in association cortices and lower variability in sensorimotor cortices (r = 0.4626, p < 0.0001; Fig. S11e). Consistent with HCP-D, the age effect on whole-brain mean vFC was not significant (Fig. S11g). Of the 39 HCP-D regions with significant age effects, 23 (59.0%) were replicated in PNC under the same significance criterion (ANOVA FDR-corrected p < 0.05; Fig. S11f). Thus, the PNC replication supported the shared cortical hierarchy of vIFP and vFC and the dissociation between a significant global age-related increase in vIFP and regionally specific maturation of vFC. Intra-subject vFC was estimated from two equal-length segments of the resting-state acquisition and regressed out of inter-subject vFC. Nevertheless, because the PNC dataset included only one resting-state session, we had to incorporate two task-based sessions into the vFC analysis. Task-evoked neural activity may therefore have affected the replication of some regional vFC effects.

## DISCUSSION

In a large developmental cohort, we delineated two complementary dimensions of inter-individual functional variability: variability in individualized functional parcellation (vIFP) and variability in functional connectivity (vFC). We also characterized their developmental patterns, multiscale structural substrates, and associations with cognitive measures. Our study demonstrated that both vIFP and vFC were anchored to the S-A axis, while vIFP showed a progressive global increase and vFC showed regionally specific age-related changes without a significant whole-brain age effect. Importantly, this maturation was non-linear, characterized by a synchronized developmental phase shift between ages 14 and 16 years, during which brain functional organization transitioned to accelerated individualization. Structural variability showed dissociable developmental trends across measures: variability in individualized white-matter structural connectivity (vSC) and individualized morphometric similarity networks (vMSN) increased with age, whereas variability in individualized myelin distribution (vMD) showed no significant whole-brain changes with age. Structural variability mainly underpinned variability in functional topography, and the coupling between these variabilities showed significant age effects across brain regions. Moreover, in cognition-related analyses, vFC demonstrated stronger associations with cognitive variability than vIFP. Taken together, the two distinct forms of functional variability displayed overlapping spatial distributions but divergent developmental profiles, structural substrates, and cognitive associations.

In the contemporary era of functional neuroimaging, brain function is often conceptualized as operating within a distributed, large-scale network framework^56,57^. Inter-individual variability in brain functional organization manifests in two principal, dissociable forms: topographic variability (vIFP) and connectional variability (vFC) ^18,25,58^. Our findings indicated that vIFP was more prominent within transmodal association cortices, a discovery that carries profound methodological implications. Specifically, neuroimaging analyses that rely on population-level atlases may suffer from topographic aliasing. In regions of high spatial diversity, these group templates implicitly incorporate topographic variance into connectivity estimates, thereby spuriously inflating inter-subject variability in vFC. By employing individualized functional parcellation, we sought to disentangle these two forms of functional variability and to examine each of them separately. Our results showed that functional connectivity derived from subject-specific, IFP-defined boundaries was significantly attenuated in the association cortex compared to atlas-based estimates, presumably reflecting a more precise representation of cortical functional architecture.

While previous studies across the adult lifespan have reported significant increases in vFC^11,59^, our findings derived from a high-precision IFP framework provide a critical refinement to this conclusion. Specifically, the prominent age-related effect in functional variability was primarily an increase in vIFP rather than vFC. Our results revealed that both vIFP and vFC converged on a pivotal inflection point between ages 14 and 16 years, signaling a sensitive developmental window of large-scale functional reorganization with personalization and fine-tuning of neural architecture^4,60,61^. Our results are consistent with this view, indicating that the mid-adolescent inflection may index a transition from relatively conserved functional organization toward more rapid individuation of higher-order functional topography^12,62^. Moreover, as shown in Fig. 3e, f, developmental changes in vIFP appeared to be driven mainly by inter-individual variability in the position rather than the size of IFPs, implying that relocation and boundary refinement of functional regions are key to the topological personalization of functional organization in youth. In contrast, intra-individual variability in vFC followed an opposite trajectory, declining across most regions (Fig. S4b, c). This pattern likely reflected maturation of the nervous system toward more stable, coherent, and efficient intra-individual functional communication. The combination of increasing inter-individual heterogeneity in higher-order networks and simultaneous intra-individual stabilization paints a nuanced picture of adolescent brain development: on the one hand, brain organization becomes more stable and integrated within individuals; on the other hand, higher-order functional organization shows greater inter-individual differentiation. Collectively, these dual dynamics suggest that the brain balances biological stability for processing efficiency with topographic uniqueness to support the adaptive flexibility required for complex cognition.

A key contribution of this work is the systematic elucidation of the multiscale structural substrates that underpin individualized functional organization. By quantifying vSC, vMSN, and vMD within the high-precision reference frame of IFP, our study revealed their distinct spatial patterns and developmental trajectories. Structural variability also adhered to the S-A axis, with greater inter-individual variability in association cortices. Notably, the developmental trajectories of these structural metrics were largely dissociated. While vSC and vMSN showed progressive individualization at the whole-brain level, vMD remained remarkably stable across youth. This structural foundation ensures that neural communication remains consistently efficient during brain maturation, despite substantial inter-individual differences in the topology of functional organization^63,64^. In contrast, the expansion of vMSN likely reflected the protracted individualization driven by synaptic pruning and experience-dependent remodeling, contributing to the modularization and individualization of cortical architecture, especially in higher-order regions^65,66^. These structural metrics also highlight a fundamental balance between stable efficiency and individualized flexibility during neurodevelopment. The trajectories of structural and functional variability point to a shared developmental inflection between 14 and 16 years of age. This mid-adolescent transition appears closely linked to the hierarchical sequence of cortical maturation. At approximately this age, intrinsic-activity development reached maximal alignment with the S-A axis^61^. Concurrently, structural-connectivity maturation shifted from sensorimotor-dominant to association-dominant strengthening^67^. Furthermore, ROI-level coupling analyses revealed that structural variability preferentially constrained vIFP rather than vFC. This implies that individual variability in brain anatomy (white matter connectivity patterns, myelin, and cortical morphology) exerts a more direct influence on the spatial topography of functional organization. The structure–function coupling exhibited significant age effects throughout youth, with especially pronounced coupling in the default mode, attention, and limbic networks, underscoring the complex, synchronized structural and functional plasticity of these higher-order networks during development. The primary trajectory of coupling strength was characterized by a widespread decline. This reduction was particularly pronounced within the networks of the association cortex. The “tethering hypothesis” provides a compelling evolutionary framework for these observations. It posits that the disproportionate expansion of the human association cortex during evolution drove the emergence of both advanced cognitive capacities and remarkable inter-individual variability^68^. The developmental decoupling between structure and function represents a progressive release of functional organization from its underlying structural tethers. This dynamic untethering ultimately steers the cortex toward a highly individualized functional architecture.

However, when linking functional variability to cognitive abilities (e.g., fluid and crystallized intelligence), we observed a nuanced pattern of distinction and complementarity between the two primary functional dimensions. Although vIFP captured inter-individual variability in spatial topography, its predictive power for cognition was markedly weaker than that of vFC. Furthermore, integrating vIFP with vFC did not reliably enhance predictive accuracy. This finding prompted a critical reflection: despite substantial inter-individual variability in the spatial location and shape of functional regions, homologous functional regions may perform equivalent roles or subserve similar functions across individuals. Consequently, the conservation of core functional attributes may dampen the predictive utility of vIFP for cognitive variability. In contrast, vFC showed robust predictive performance for inter-individual cognitive variability, particularly crystallized intelligence. This supports the view that, while the spatial topology of vIFP constitutes an important, structure-associated feature of individuality, variability in connectivity patterns (vFC) may more directly reflect information-processing efficiency and the uniqueness of cognitive function. Together, these findings highlight the need to jointly consider intra-regional organization (topography) and inter-regional interactions (connectivity) to understand how the personalized human brain processes information and generates complex behavior during the transition into adulthood^18,69^.

Several limitations should be acknowledged. First, while the HCP-D provides a high-quality sample, it is predominantly cross-sectional, which may underestimate true longitudinal developmental trends. This is particularly relevant for conclusions regarding the coupling between functional variability and structural variability. Future longitudinal studies may further leverage within-subject repeated measurements to examine whether structure and function mutually constrain each other over development and to move beyond association toward causal inference. Second, our IFP framework was initialized from a population atlas and optimized for functional homology, which led us to retain only the 87 parcels that were identifiable across all participants (87/116). Importantly, the excluded parcels likely reflect meaningful inter-individual variability precisely because they are not universally present (or not consistently detectable) across individuals. Consequently, our estimates of functional variability are restricted to universal parcels, and the developmental profile and behavioral relevance of functional variability in these non-universal regions remain open questions for future work. Third, most current individualized functional parcellation approaches require multiple repeated scans per participant to obtain sufficiently long, high-quality fMRI time series, which constrains feasibility for large-sample, wide age-range developmental studies. Methodological advances are needed to derive accurate individualized parcellations from shorter acquisitions, enabling scalable individualized mapping across the lifespan.

In summary, this work established a multidimensional framework that delineates topographic variability (vIFP) and connectional variability (vFC) as dissociable yet complementary facets of brain functional organization. While both functional indices exhibited convergent spatial distribution patterns along the S-A axis and converged on a pivotal maturational inflection point between ages 14 and 16 years, they embodied a divergence in their structural constraints and cognitive significance. vFC emerged as a significantly stronger neural correlate of cognitive variability, whereas vIFP appeared to serve as a stable, structurally anchored feature of individual identity. The dual-perspective framework enriched our understanding of the personalization of brain development in youth. Future research should extend these findings to investigate the roles of functional and structural variability in neurodevelopmental disorders and examine how environmental and genetic factors jointly shape this individual variability, thereby informing more precise diagnostic and intervention strategies.

## METHODS

### Participants

Data for the discovery analyses were obtained from HCP-D Release 2.0. We initially selected 627 participants (339 females; mean age = 14.50 ± 4.03 years; age range = 5.58-21.92 years) for data-completeness screening. After quality control, the final HCP-D discovery cohort comprised 601 participants aged 6–22 years. The original collection of HCP-D data was approved by the Washington University Institutional Review Board (IRB No. 201204036). In accordance with the Declaration of Helsinki, written informed consent was obtained from adult participants and from the parents or legal guardians of minor participants.

MRI data for the HCP-D were acquired on a Siemens 3 T Prisma scanner, with detailed parameters reported in Ref^52^. Structural imaging included T1-weighted (T1w) and T2-weighted (T2w) sequences: the T1w images were obtained using a 3D multi-echo MPRAGE sequence (0.8 mm isotropic voxels; TR/TI = 2500/1000 ms; TE = 1.8/3.6/5.4/7.2 ms; flip angle = 8°), whereas the T2w images were acquired using a 3D variable-flip-angle turbo-spin-echo (TSE) SPACE sequence (0.8 mm isotropic voxels; TR/TE = 3200/564 ms). Diffusion MRI data were acquired using a 2D multiband (MB) spinlJecho echo-planar imaging (EPI) sequence (MB4, 1.5 mm isotropic voxels, 185 directions, b-values = 1500/3000 s/mm2, with 28 b = 0 s/mm2 reference volumes). Resting-state fMRI was acquired using a 2D multiband gradient-recalled echo (GRE) EPI sequence (MB8, 2.0 mm isotropic voxels, TR/TE = 800/37 ms, flip angle = 52°). Participants aged 8–21 years underwent 26 minutes of rs-fMRI scanning across four runs (6.5 min/run), whereas the 5–7-year cohort completed 21 minutes through six shortened runs (3.5 min/run)^51,52,70^.

To evaluate the generalizability of the findings from the HCP-D discovery cohort, we analyzed an independent sample from the Philadelphia Neurodevelopmental Cohort (PNC). The PNC dataset included 1,375 participants. After quality control, participants were further required to have all 87 homologous functional regions defined in the HCP-D dataset, yielding a final replication sample of 610 participants aged 8–23 years (mean age = 16.35 ± 3.13 years; 346 females). The original collection of PNC data was approved by the Institutional Review Boards of the University of Pennsylvania and the Children’s Hospital of Philadelphia (IRB-approved protocol number 810336). In accordance with the Declaration of Helsinki, written informed consent was obtained from participants aged 18 years or older, and written assent and parental/legal guardian permission were obtained for minors.

All MRI data were acquired using the same Siemens 3 T Tim Trio whole-body scanner equipped with a 32-channel head coil at the Hospital of the University of Pennsylvania. T1-weighted images were acquired using a 5-min magnetization-prepared rapid gradient-echo sequence (MPRAGE; TR/TE/TI = 1810/3.51/1100 ms; field of view = 180 × 240 mm²; matrix = 192 × 256; voxel size = 0.9 × 0.9 × 1 mm³). Functional imaging included one resting-state scan and two task-based scans involving n-back working memory and emotion identification. All fMRI data were acquired using the same single-shot, interleaved multislice gradient-echo echo-planar imaging sequence (TR/TE = 3000/32 ms; flip angle = 90°; field of view = 192 × 192 mm²; matrix = 64 × 64; 46 slices; slice thickness/gap = 3/0 mm; voxel size = 3 mm isotropic). The resting-state, n-back, and emotion-identification scans contained 124, 231, and 210 volumes, respectively, providing approximately 28 min of fMRI data in total^7,53^.

### Preprocessing

All structural and functional images underwent HCP minimal processing 60. Briefly, structural T1w and T2w images underwent gradient distortion correction, alignment, bias field correction, registration to Montreal Neurological Institute (MNI) space, white-matter and pial surface reconstruction, structural segmentation, and surface registration and downsampling to the 32k_fs_LR mesh. Regarding fMRI data, the preprocessing pipeline included spatial distortion correction, motion correction, EPI distortion correction, registration to MNI space, intensity normalization, mapping of volume time series to 32k_fs_LR mesh, and smoothing with a 2-mm surface kernel. For diffusion MRI, we employed the standard preprocessing steps implemented in MRtrix3. Diffusion images were denoised, Gibbs ringing artifacts were removed, and eddy current-induced distortions, head motion, signal dropout, EPI distortion, and B1 field inhomogeneity were corrected using MRtrix3.

For the PNC replication cohort, T1-weighted images and all three fMRI acquisitions were preprocessed using DeepPrep v25.1.0^72^. This workflow independently generated cortical reconstructions, preprocessed BOLD data, and quality-control outputs without using existing fMRIPrep derivatives. The resting-state, n-back, and emotion-identification scans were processed using the same workflow. DeepPrep generated volumetric BOLD outputs in MNI space at 2-mm resolution and surface BOLD time series on fsaverage6. For compatibility with the individualized functional parcellation procedure, the fsaverage6 time series were resampled to fsaverage4 using FreeSurfer’s mri_surf2surf. Resampling was performed separately for each hemisphere, with fsaverage6 and fsaverage4 specified as the source and target surfaces, respectively. This reduced the surface resolution from 40,962 to 2,562 vertices per hemisphere. Because both templates are FreeSurfer standard surfaces, no intermediate volumetric transformation was applied. The resulting fsaverage4 time series from all three acquisitions were used in the PNC replication analyses.

### Quality Control

We applied a multi-step quality control procedure combining automated image-quality assessment, processing checks, and expert visual review. T1w, T2w, and fMRI scans were first assessed using MRIQC, and scans were excluded if they were flagged as outliers on three or more modality-specific no-reference quality metrics. Images that failed preprocessing or downstream processing were also excluded. Structural data were further screened using the Euler number to assess cortical surface reconstruction quality. Functional runs were excluded for excessive head motion (mean FD > 0.5 mm or >20% of frames with FD > 0.5 mm), fewer than 100 retained time points, or a retained-to-original time-point ratio <90%.

All remaining data underwent visual QC by four trained reviewers. Structural images were inspected for artifacts, cortical segmentation, surface reconstruction and registration quality; T1w/T2w-derived myelin maps were inspected for abnormal distributions; and fMRI data were assessed for brain coverage, registration quality and volume-to-surface mapping. Participants with any unresolved quality defect were excluded.

### Individualized Functional Parcellation

We identified homologous functional regions in individual subjects using an iterative segmentation and group-matching approach. The process began with the construction of a population-level functional atlas. This atlas was based on resting-state fMRI data from 1,000 healthy adults and initially divided the cerebral cortex into 17 large-scale functional networks, following Yeo et al. Building on this foundation, the hand motor area was isolated. Subsequently, based on the continuity of regional locations on the cortex, these networks were further subdivided to form a final population-level atlas comprising 116 functional regions corresponding to 18 functional networks.

For individual subjects, BOLD time series data were resampled to the fsaverage4 template space, resulting in 4,771 cortical vertices across the whole brain, excluding the corpus callosum. Initially, the 18 functional networks from the population-level atlas were applied to each individual. For each network, the time series of all vertices within it were averaged to derive its ‘core signal’. Subsequently, each vertex’s time series was correlated with these 18 core signals, and vertices were reassigned to the network with which they exhibited the highest correlation. This reassignment process was iterated: core signals were recalculated based on the updated vertex assignments, and vertices were reassigned again. This iterative, individual-data-driven updating of vertex network labels progressively reduced the influence of the population-level atlas until the assignments stabilized, thereby enabling data-driven parcellation.

Next, a clustering algorithm (using the ‘metric-find-clusters’ command in Connectome Workbench) partitioned these individual vertex-based assignments into multiple distinct patches. These individual patches were then matched to the 116 functional regions of the population-level atlas. This matching procedure was based on the hypothesis that ROIs within the population-level atlas roughly represent the centroids of homologous ROIs from different individuals. The matching was primarily performed by assessing vertex overlap and geodesic distance between individual ROIs and atlas ROIs. Any individual ROI that did not overlap with any atlas ROI, or whose geodesic distance exceeded a predefined threshold, was marked as ‘unidentified’. Finally, the homologous functional parcellations were defined as those ROIs that were successfully identified across all subjects. In most cases, the number of resulting homologous functional parcellations was fewer than 116.

### Inter-individual Variability

Inter-individual variability for each neuroimaging feature was quantified using individualized functional parcellations (IFPs). For each feature, variability was quantified between every pair of subjects. Group-level inter-individual variability was calculated as the average of pairwise inter-subject variability within the group.

#### (1) Functional Parcellation (vIFP)

Individualized functional parcellations (IFPs) were obtained for each subject in the fsaverage4 cortical space. Inter-subject similarity for the same region of interest (ROI) was quantified using the Dice coefficient, which ranges from 0 to 1. We used 1 - Dice to represent inter-subject variability in functional parcellation (vIFP).

#### (2) Position (vPosition)

Compared to some existing parcellation methods, our individualized functional parcellation approach was designed to maintain discrete ROIs on the cortex while ensuring that individual ROIs encompassed contiguous cortical areas. Consequently, we used the centroid of each ROI to represent its spatial location. Inter-subject variability in IFP position (vPosition) was defined as the Euclidean distance between the centroids of the same ROI across different individuals.

#### (3) Area (vArea)

Individual functional parcellation labels in the fsaverage4 cortical space were registered to individual subjects’ native cortical spaces using FreeSurfer’s cortical registration tool (mri_surf2surf). Using FreeSurfer’s anatomical statistics tool (mris_anatomical_stats), the surface area of each ROI was calculated. Inter-subject variability in functional parcellation area was defined as the absolute difference in surface area for the same ROI between different individuals.

#### (4) Functional Connectivity (vFC)

Individualized functional connectivity (FC) matrices for each subject were constructed based on the individualized functional parcellations. As described previously, BOLD time series were resampled to the fsaverage4 template space, yielding 4,771 whole-brain vertices, excluding the corpus callosum. Concurrently, the individualized brain functional parcellations provided the partition labels in the fsaverage4 template space. The BOLD signals of all vertices within each ROI were averaged to serve as the ‘core signal’ for that ROI. FC matrices were constructed using the core signals of each ROI. FC between two ROIs was defined as the Pearson correlation coefficient between their core signals. A total of 87 functionally homologous regions were identified, resulting in an 87 × 87 FC matrix, which was then Fisher’s Z-transformed. For subjects with multiple sessions, FC matrices were constructed using data from each session and then averaged to obtain the subject’s final FC matrix. The calculation workflow for inter-subject variability in FC is illustrated in Fig. S10.

Each row in the FC matrix represents an ROI’s FC profile, describing its functional connectivity architecture with all other ROIs on the cortex. Inter-subject variability in FC was estimated through the inter-subject variability of these FC profiles. For a specific ROI, inter-subject FC similarity was quantified as the Pearson correlation between the corresponding FC profiles, as shown in Equation (1):

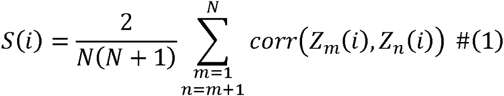

Here, *Z_m_*(*i*) and *Z_n_*(*i*) represent the FC profile vector for the ROI *i* of subjects *m* and *n*, respectively. *corr*(*Z_m_*(*i*), *Z_n_*(*i*)) denotes the FC similarity for the ROI *i* between these two subjects. Within a group of subjects, pairwise FC similarities for all subjects were computed and averaged, denoted as *S*(*i*). The inter-subject variability for the ROI *i* in this group was then expressed as *V_intra_*(*i*), according to Equation (2):

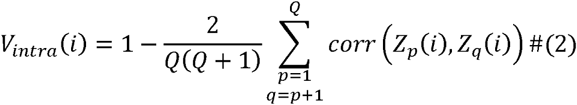

The intra-subject variance was then regressed out using ordinary least-squares regression (i.e., a general linear model, GLM). The inter-subject variability for the ROI *i* in this group was then expressed as *V_intra_*(*i*), according to Equation (3):

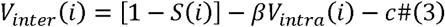

Here, β and *c* are parameters determined via ordinary least-squares.

#### (5) Structural Connectivity (vSC)

Structural connectivity (SC) was reconstructed using probabilistic tractography implemented in MRtrix3, grounded in individualized functional parcellation. Processing began by extracting the mean b=0 image from the bias-field-corrected DWI data using MRtrix3’s dwiextract -bzero command, followed by averaging across volumes via mrmath -axis 3 mean. The b0 image was registered to the T1w image using ANTs’ antsRegistrationSyN.sh script to compute an affine transformation matrix. A DWI-space binary mask was generated using MRtrix3’s dwi2mask to enhance the biological plausibility of streamlines and optimize computational efficiency. Tissue-specific response functions for white matter (WM), gray matter (GM), and cerebrospinal fluid (CSF) were estimated using the dhollander method (dwi2response dhollander). Multi-shell multi-tissue constrained spherical deconvolution (MSMT-CSD) was subsequently performed with dwi2fod msmt_csd to compute fiber orientation distributions (FODs). Global intensity normalization was applied using mtnormalise. A five-tissue-type (5TT) image was generated using 5ttgen freesurfer, from which the gray-white matter interface (GMWI) was extracted via 5tt2gmwmi. Whole-brain probabilistic tractography was implemented in tckgen using anatomically constrained tractography (ACT) with the 5TT image provided via the -act flag. Streamlines were seeded from the gray-white matter interface (-seed_gmwmi) and reconstructed using the second-order integration over FODs (iFOD2) algorithm (-algorithm iFOD2). A total of 5 million streamlines were initially generated (-select 5000000), followed by spherical deconvolution-informed filtering (SIFT) with tcksift to mitigate length-related biases, retaining 1 million biologically plausible streamlines (-term_number 1000000). The individualized functional parcellation labels were co-registered to DWI space using ANTs’ antsApplyTransforms, guided by the precomputed affine matrix, for downstream SC matrix construction. A symmetric 87 × 87 SC matrix was constructed using MRtrix3’s tck2connectome, representing streamline counts between functional regions. Edge weights were scaled by the inverse product of region volumes (-scale_invnodevol) to correct for region size heterogeneity. The matrix diagonal was set to zero (-zero_diagonal), and symmetry was enforced (-symmetric).

Based on SC matrices, we calculated inter-subject variability in SC using the same methodology as for FC.

#### (6) Morphometric Similarity Networks (vMSN)

Using FreeSurfer and the individualized functional parcellation labels, we characterized regional cortical morphology. Specifically, the individualized functional parcellation labels were registered to individual subjects’ native cortical spaces via surface registration (mri_surf2surf). Subsequently, the anatomical statistics tool (mris_anatomical_stats) was used to compute nine anatomical properties for each brain region: Number of vertices, Surface area (mm^2^), Gray matter volume (mm^3^), Average thickness (mm), Thickness standard deviation (mm), Integrated rectified mean curvature (1/mm), Integrated rectified gaussian curvature (1/mm^2^), Folding index, and intrinsic curvature index. Because of the high collinearity between ‘Number of vertices’ and ‘Surface area (mm^2^)’, ‘Number of vertices’ was removed. Ultimately, the structural morphology feature vector for each brain region comprised eight features.

Inter-subject variability in regional structural morphology was defined as the difference between these structural morphology feature vectors. The similarity in structural morphology for the same brain region between two individuals was represented by the Pearson correlation coefficient (r) of their corresponding structural morphology feature vectors. Inter-subject variability was then defined as the complement of the correlation (1 − *r*). Within a group of subjects, pairwise differences were calculated and then averaged.

#### (7) Myelin Distribution (vMD)

Myelination was assessed using the T1w/T2w ratio, which indexes intracortical myelin and was produced for each participant by the HCP minimal pipeline. Subsequently, individual myelin distribution maps were registered to the fsaverage4 surface using FreeSurfer’s surface registration tools. Using the previously obtained individualized functional parcellation labels, the T1w/T2w ratios for vertices within each functional region were extracted for individual subjects.

To compare inter-subject variability in T1w/T2w ratios within functional regions, we accounted for the fact that the number of vertices included in a functional region could vary across individuals. Accordingly, we employed the H statistic from the Kruskal-Wallis test to quantify inter-subject variability.

### Age-related Effects

To systematically investigate the age-related developmental trajectories of brain functional and structural variability during youth, we conducted analyses of age effects for various inter-individual variability measures, including variability in individualized functional parcellation (vIFP), variability in functional connectivity (vFC), variability in white-matter structural connectivity (vSC), variability in morphometric similarity networks (vMSN), and variability in myelin distribution (vMD).

Given that these measures were derived from a group of subjects, we first employed an age-sliding window approach across subjects to generate data points for subsequent statistical modeling. Specifically, all subjects were sorted by age in ascending order. A sliding window of 20 subjects was then applied with a step size of 10 subjects. For each window, we calculated the mean age of the 20 subjects within it, which served as the representative age for that window. Concurrently, all inter-individual variability measures for the subjects within that window were computed. This approach transformed the continuous age axis into a sequence of discrete data points, each representing inter-individual variability at a specific age.

Subsequently, to capture the complex non-linear relationships between these variability measures and age, we used generalized additive models (GAMs). For each inter-individual variability measure obtained through the sliding window approach (including whole-brain average values and regional values for each IFP-defined cortical region), we constructed the following GAM:

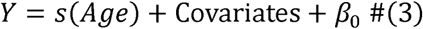

Here, *Y* represents the analyzed variability measure, *s*(*Aye*) denotes a smoothing function of age, allowing the model to capture non-linear effects, and *β*_0_ is the intercept term. For modeling vIFP, vFC, vSC, vMSN, and vMD, we included sex and head motion (mean framewise displacement, FD) as covariates.

To test the significance of age effects, we constructed two nested GAMs: one containing the age smoothing term (i.e., the model described above), and another without the age smoothing term (including only covariates and the intercept term). The two models were subsequently compared using ANOVA. The change in their residual sum of squares (or deviance) was used to calculate the F-statistic and corresponding p-value, thereby determining whether the inclusion of the age smoothing term significantly improved the model’s explanatory power for the dependent variable. By analyzing the smoothed curves obtained from the GAMs, we identified significant age effects on variability at both whole-brain and regional levels. Further analysis of the curve shapes and their first derivatives allowed us to investigate changes in variability growth rates and pinpoint critical developmental inflection points. Regarding the driving factors of vIFP, we independently analyzed the age-related changes in functional parcellation position variability (vPosition) and area variability (vArea), both obtained via the sliding window method, to compare their relative contributions to the age effects observed in vIFP.

All statistical analyses were performed in R, and p-values for regional-level significance were corrected for multiple comparisons using the false discovery rate (FDR).

### Sensitivity Analysis of Sliding-window Parameterization

The 20-subject window with a 10-subject step was used for the primary age-effect analysis. To test whether the results depended on this parameterization, we repeated the workflow with a 10-subject window and a 5-subject step. Subjects were sorted by age, and the mean age of each window represented that window. Inter-individual variability measures were recomputed within each window before GAM fitting. The sensitivity analysis used the same GAM specification, nested ANOVA test, delta R² calculation, and FDR correction as the primary analysis. Covariate handling followed the corresponding primary models, including sex and mean FD in the motion-adjusted analyses.

### Coupling of Inter-individual Variability across Features

To investigate coupling between functional variability (vIFP and vFC) and structural variability (vSC, vMSN, and vMD), as well as its age-related trajectories, we analyzed six functional–structural combinations: vIFP-vSC, vIFP-vMSN, vIFP-vMD, vFC-vSC, vFC-vMSN, and vFC-vMD. This analysis aimed to quantify the extent to which individual variability at the structural level aligned with that at the functional level, thereby revealing how function and structure co-develop at the individual scale.

For each cortical region defined by individualized functional parcellation (IFP) and for each functional-structural combination, we first calculated its inter-individual coupling strength. Specifically, for each age window (group), region, and functional-structural combination, we computed pairwise functional variability values and the corresponding pairwise structural variability values for all unique subject pairs (e.g., *vlFP_ij_* and *vlSC_ij_*, from *subject_i_* and *subject*_j_). These pairwise functional values were then aggregated into a functional variability vector, and the corresponding structural values were aggregated into a structural variability vector. Both vectors had a length equal to the number of unique subject pairs. Finally, we computed the Pearson correlation coefficient between these two variability vectors. The resulting coefficient represented the inter-individual coupling strength for that specific region, functional–structural combination, and age window.

Using these steps, a coupling strength value was obtained for each region, each functional–structural combination, and each age window. To investigate whether these coupling strengths changed significantly with age, we used generalized additive models (GAMs). The calculated coupling strengths were used as the dependent variable and the mean age of the window as the independent variable, with a smoothing function incorporated to capture non-linear relationships. By testing the significance of the smooth term for age, we identified the developmental trajectories of inter-individual functional-structural coupling during youth.

### Prediction of Cognitive Variability

To thoroughly investigate how individual variability in brain functional organization predicts cognitive variability during youth, we conducted a series of subject-pair-based prediction analyses. These analyses aimed to identify brain functional variability features significantly associated with inter-individual variability in specific cognitive domains (crystallized intelligence, fluid intelligence, and total intelligence). Unlike the aforementioned analyses focusing on group variability or age trajectories, these analyses directly predicted cognitive differences between any two subjects, without using age-sliding windows or age-effect analyses.

The cognitive measures used included crystallized intelligence, fluid intelligence, and total intelligence, all derived from the Human Connectome Project – Development (HCP-D) dataset and measured using the NIH Toolbox. For each cognitive measure, we calculated the absolute difference in cognitive scores between all unique subject pairs, which served as the target cognitive variability to be predicted.

As predictors, we constructed functional inter-individual variability features for each subject pair. Specifically, for every pair of subjects, we extracted two types of brain functional variability features: vIFP and vFC. These variability features constituted the independent variables in our prediction models.

We employed ridge regression, which is particularly suitable for handling high-dimensional brain feature data that may exhibit multicollinearity. By applying an L2-norm penalty to the regression coefficients, ridge regression prevents model overfitting and enhances generalization performance. We constructed three independent prediction models to assess the relative contributions of different types of functional variability to cognitive variability:

1. vIFP-only variability prediction model: used only the vIFP features between subject pairs to predict cognitive variability.
2. vFC-only variability prediction model: used only the vFC features between subject pairs to predict cognitive variability.
3. Combined vIFP and vFC variability prediction model: used both vIFP and vFC features between subject pairs to predict cognitive variability.

To evaluate model performance, all prediction models were trained and assessed using 100 repetitions of 2-fold cross-validation. In each cross-validation fold, the entire subject-pair dataset was randomly partitioned into training and test sets. To ensure balance in key demographic characteristics between the training and test sets, a stratified sampling strategy was employed: subjects were first divided into five age strata, then further stratified by sex within each age stratum. Subsequently, subjects were randomly divided into two halves, forming two mutually exclusive subject-pair subsets for training and testing. Prediction accuracy was expressed as the Pearson correlation coefficient (*r*) between the predicted and actual cognitive difference values on the test set. To assess the statistical significance of prediction accuracy, we performed 100,000 permutation tests, constructing a null distribution by randomly shuffling cognitive variability labels and calculating the empirical p-value for the observed Pearson *r*. All prediction analyses were completed in MATLAB.

## Acknowledgments

The study was supported by the Scientific and Technological Innovation (STI) 2030 Major Projects 2021ZD0200500 (https://en.most.gov.cn/), the National Natural Science Foundation of China (https://www.nsfc.gov.cn/english/site_1/index.html, 32271146 to S.L.), and Startup Funds for Top-notch Talents at Beijing Normal University.

## Author contributions

Conceptualization: SYL, ZKY

Methodology: ZKY, DBZ, XXD

Investigation: ZKY

Visualization: ZKY, YRH, JCZ

Supervision: SYL

Writing—original draft: ZKY

Writing—review & editing: SYL, XXD, YRH, DBZ, LC, ZXC, XYW

## Data and materials availability

The data used in this study are available through the Human Connectome Project–Development (HCP-D; https://www.humanconnectome.org/study/hcp-lifespan-development), a controlled-access repository, and the Philadelphia Neurodevelopmental Cohort (PNC), which is publicly available through the Reproducible Brain Charts portal (https://reprobrainchart.github.io/). The code used for the analyses will be made publicly available on GitHub upon acceptance of the manuscript.

## Code Availability

The code for this manuscript is available (https://github.com/Izaackk/youth-ifv-hfr). Software packages used in this manuscript include HFR (https://github.com/MeilingAva/Homologous-Functional-Regions)^28^, HCP pipeline v4.4.0-rc-MOD-e7a6af9 (https://github.com/Washington-University/HCPpipelines/releases), FreeSurfer v6.0.0 (https://surfer.nmr.mgh.harvard.edu/), FSL v6.0.5 (https://fsl.fmrib.ox.ac.uk/fsl/fslwiki), Connectome Workbench v1.5.0 (https://www.humanconnectome.org/software/connectome-workbench), DeepPrep 25.1.0 (https://github.com/pBFSLab/DeepPrep)^72^, MATLAB R2020b (https://www.mathworks.com/products/matlab.html), SPM12 toolbox v6470 (https://www.fil.ion.ucl.ac.uk/spm/software/spm12), cifti-matlab toolbox v2 (https://github.com/Washington-University/cifti-matlab), and R v4.4.1 (https://www.r-project.org).

