## Supplementary Information for "Spatiotemporal Patterns and Structural Substrates of Individual Functional Variability in Youth"

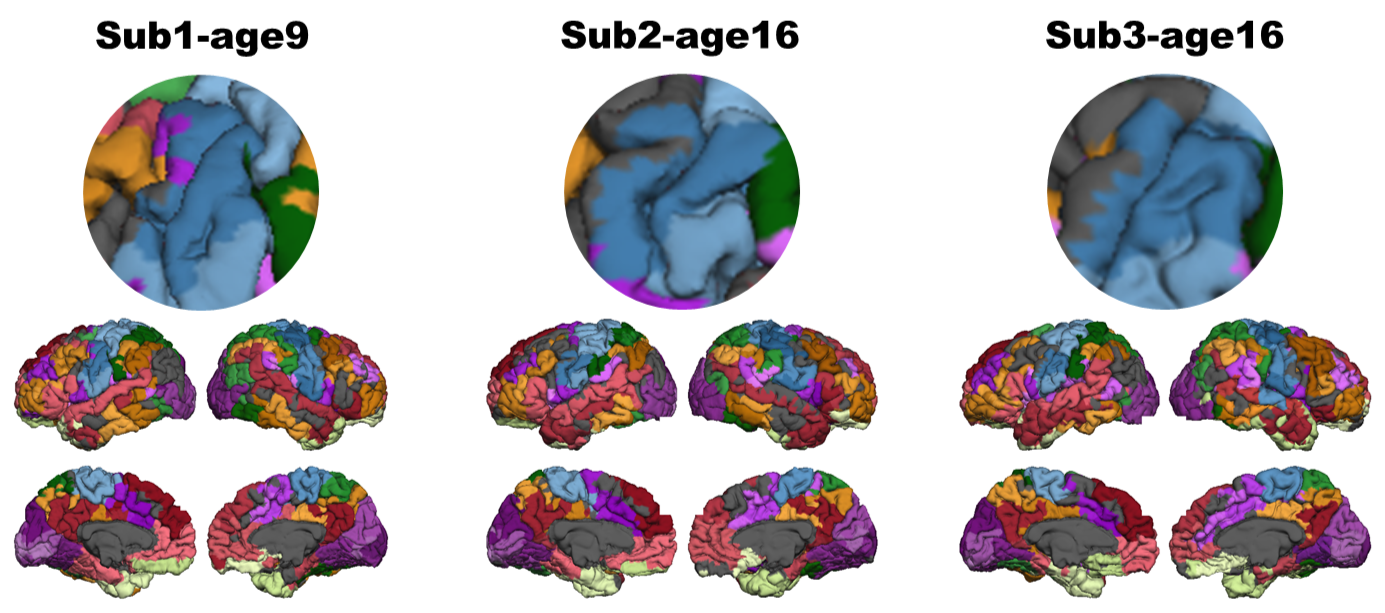

**Figure S1:** **Individualized functional parcellations of three participants of different ages.** Visualized on the pial surface, with circles indicating the face motor areas.

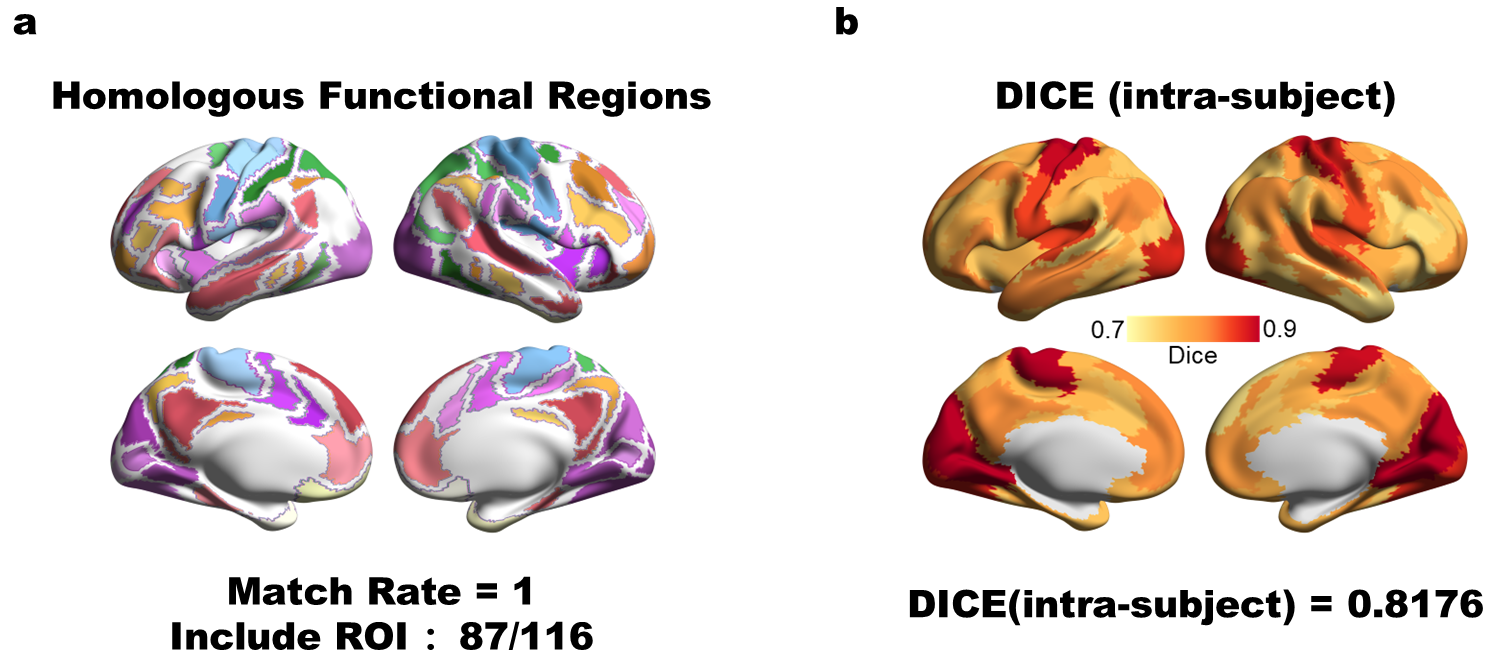

**Figure S2:** **Individualized homologous functional parcellation and its intra-individual reproducibility. a.** 87 homologous functional parcellations were identified across 601 youth participants. **b.** The intra-subject Dice coefficient of individualized functional parcellation quantifies the reliability of this method.

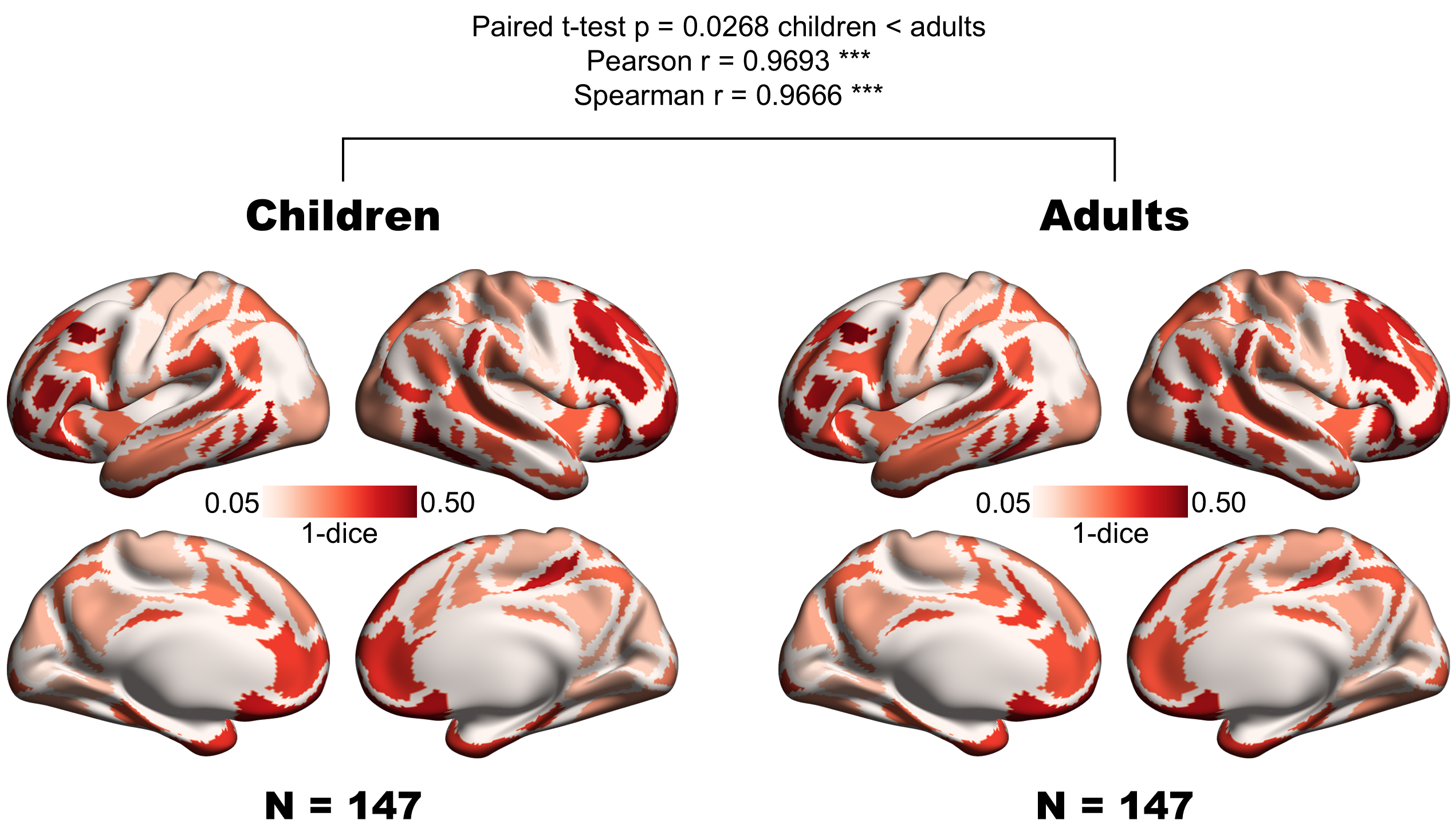

**Figure S3: Comparison of vIFP between the child and adult groups.**

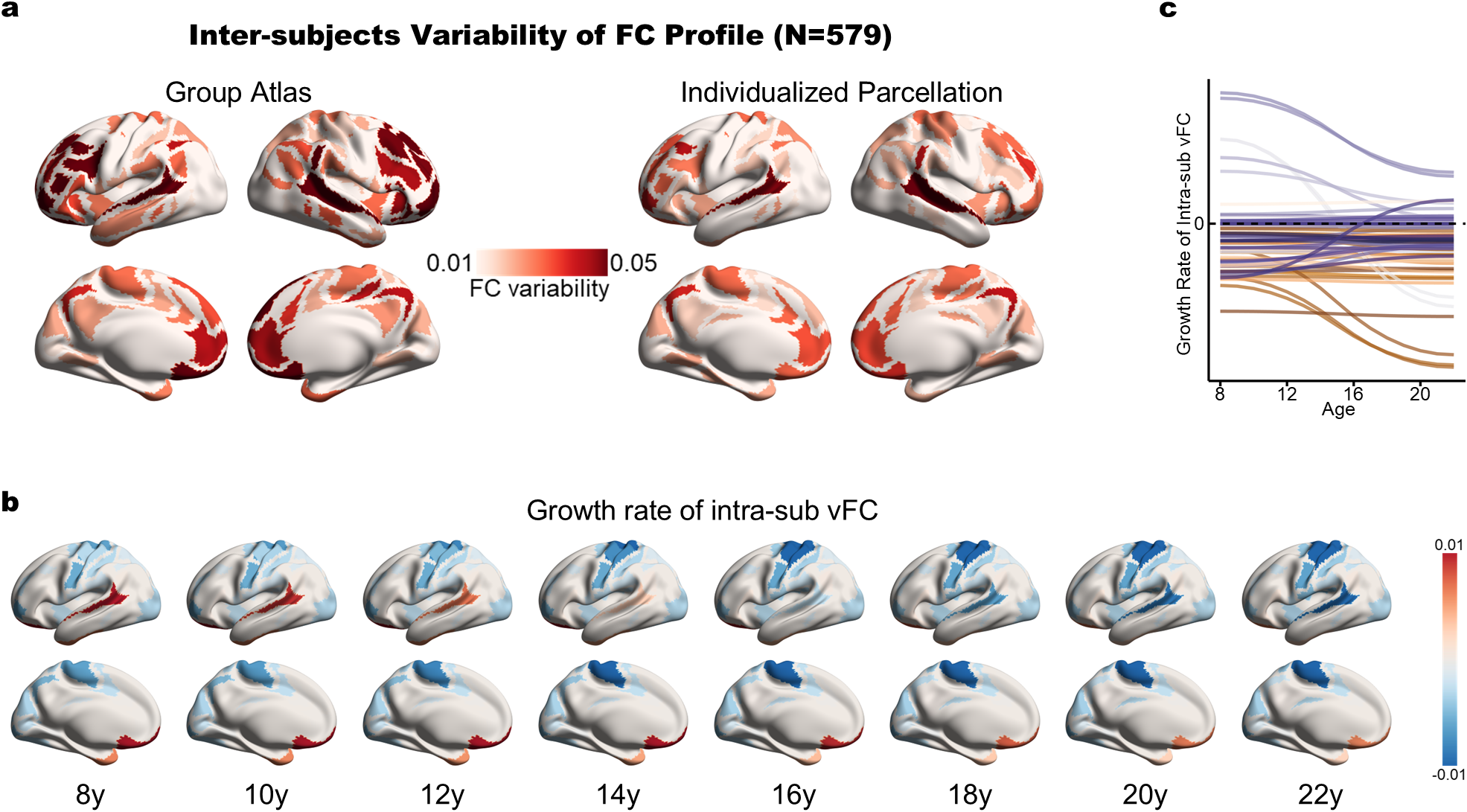

**Figure S4: Variability in FC profiles (vFC). a.** Individual variability in FC architecture was quantified in a sample of 579 participants in youth. The left panel shows results based on the population-level atlas, while the right panel shows results based on individualized functional parcellation. **b.** Growth rates of intra-subject vFC, obtained as the first derivatives of GAM-fitted developmental trajectories. **c.** Growth-rate curve for intra-subject vFC (first derivative of developmental trajectories).

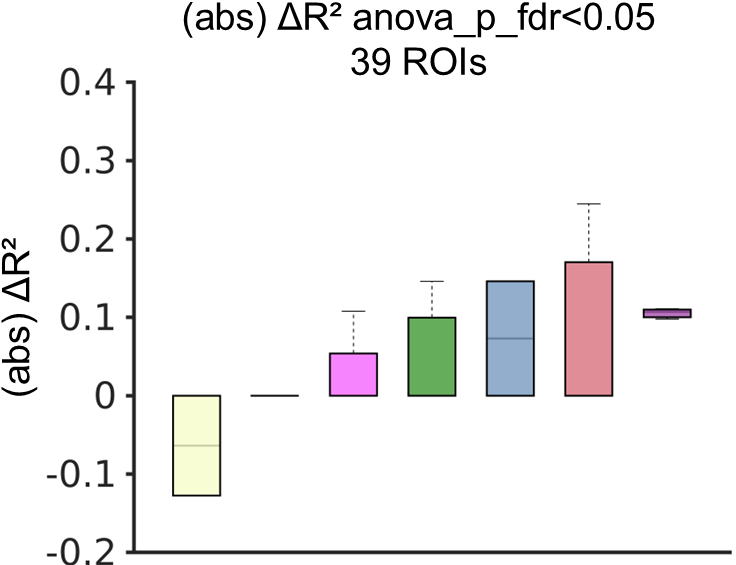

**Figure S5:** Average absolute values of delta R² for vFC across the seven networks.

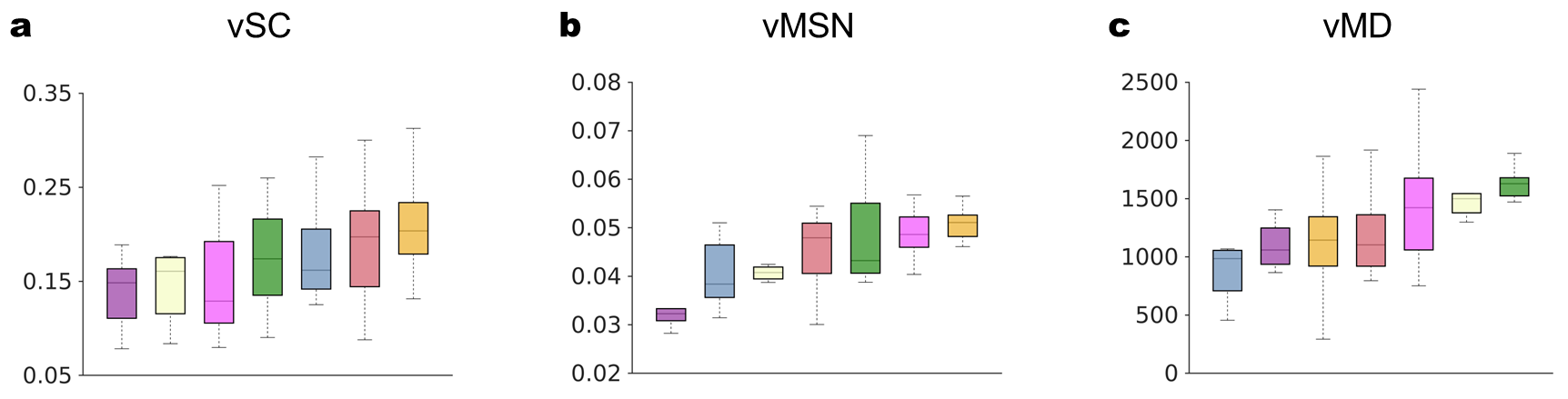

**Figure S6:** **Structural variability in 7 networks.**

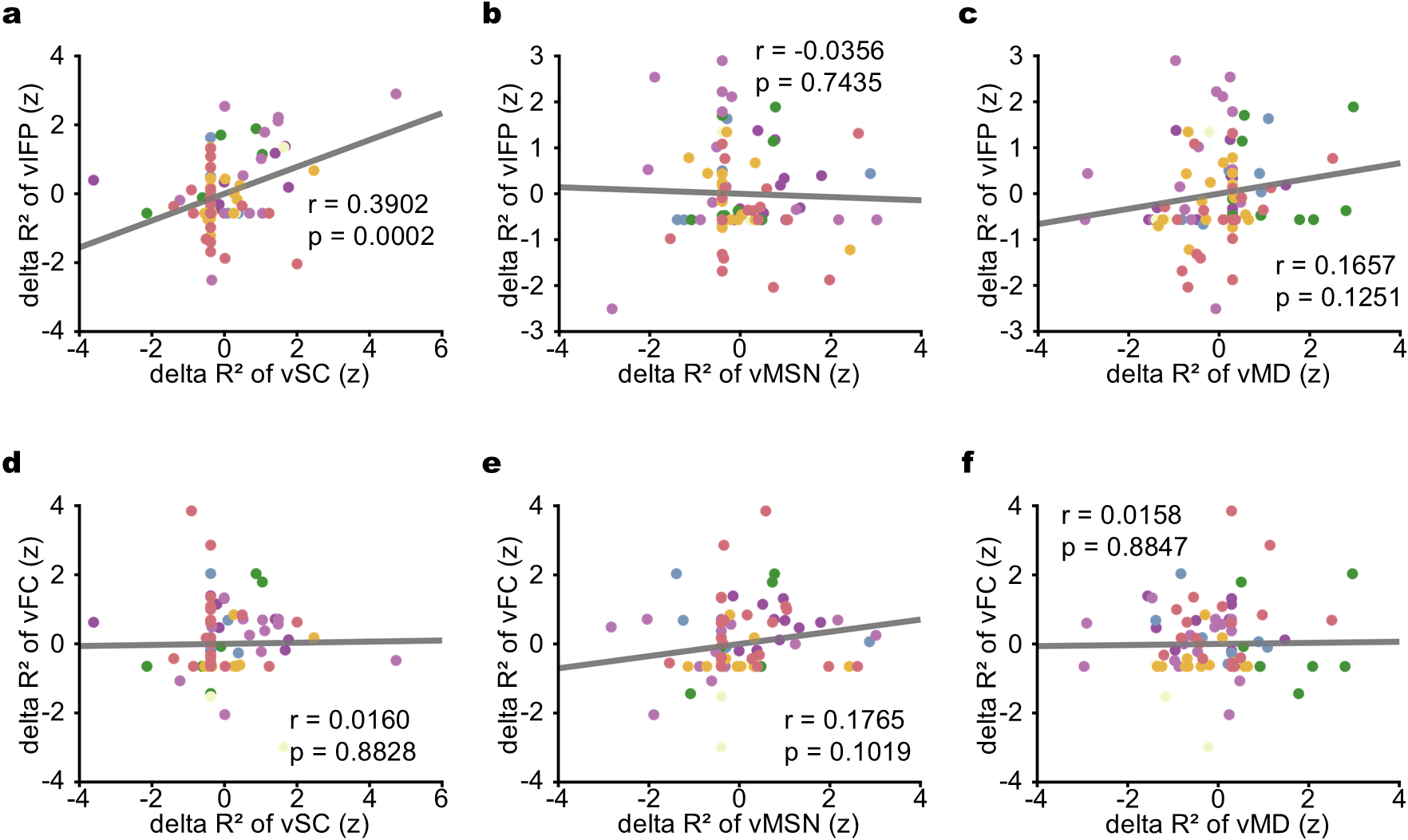

**Figure S7: Spatial similarity between the delta R² maps of functional and structural variability.**

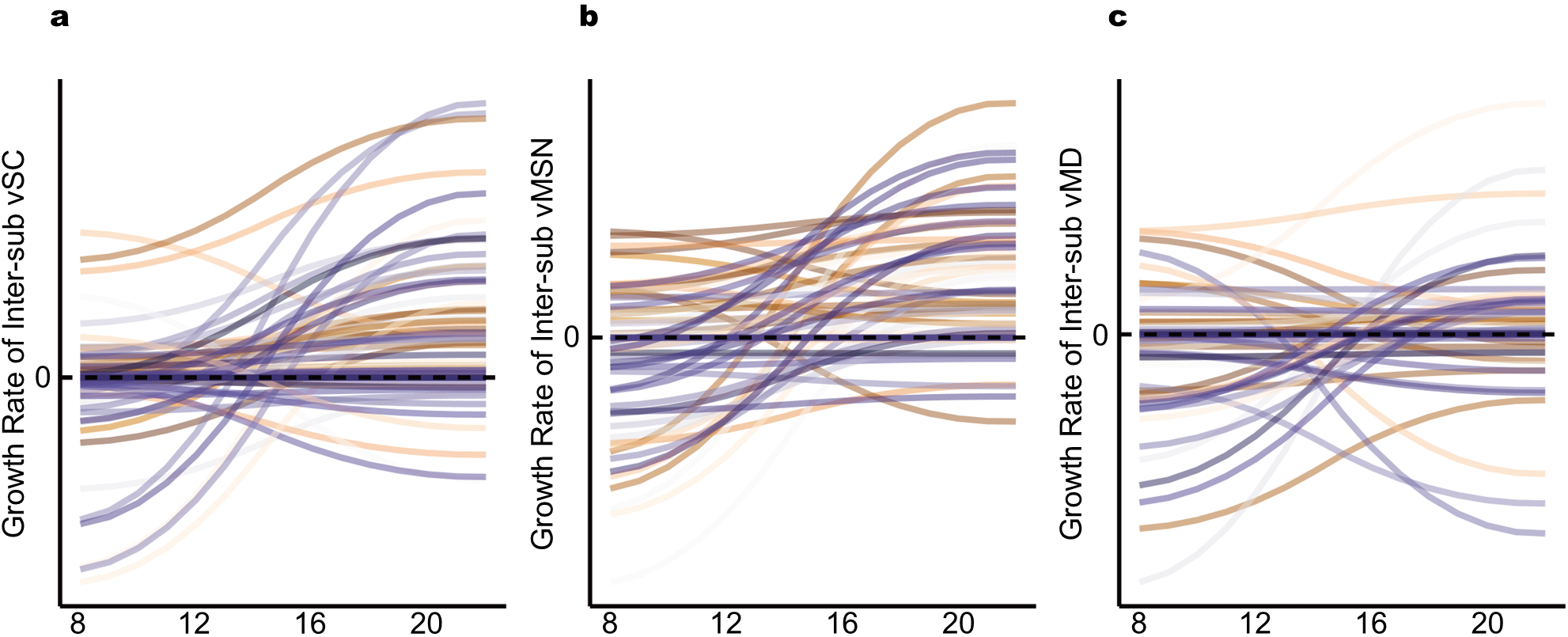

**Figure S8:** **Growth rates of structural variability. a.** Growth rate of inter-subject vSC. **b.** Growth rate of inter-subject vMSN. **c.** Growth rate of inter-subject vMD.

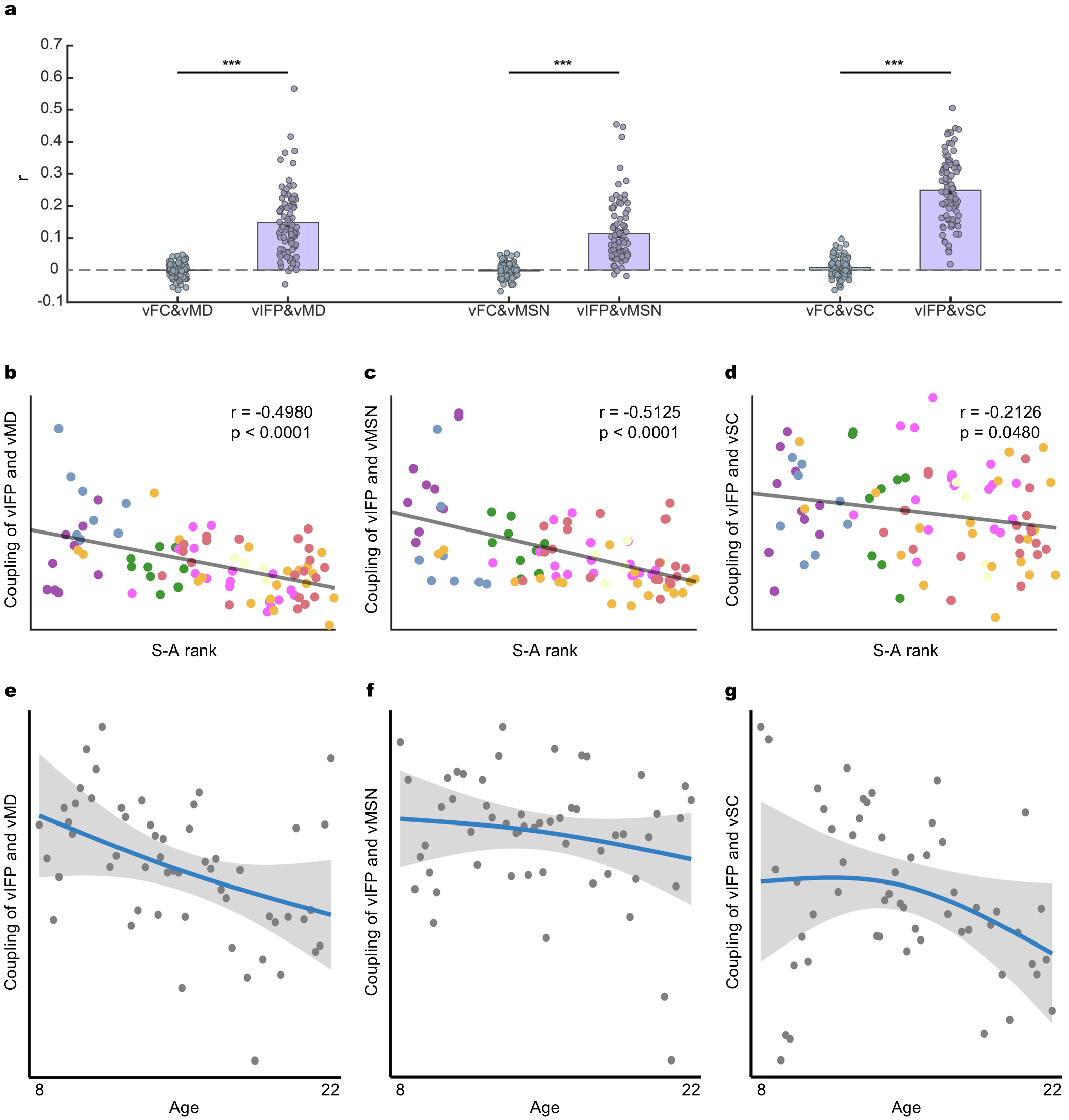

**Figure S9: Coupling between structural and functional variability. a.** Comparison of coupling strengths between two types of functional variability (vIFP and vFC) and three types of structural variability. Each scatter point represents the ROI-level coupling strength, quantified by the Pearson correlation coefficient (r), between structural and functional variability. Across all three structural phenotypes, inter-individual variability exhibited significantly stronger coupling with vIFP than with vFC. **b-d.** Spatial similarity between the sensorimotor-association (S-A) axis and the coupling strengths of vIFP with vMD, vMSN, and vSC. Each point corresponds to an ROI, color-coded according to its affiliated functional network. The coupling maps between vIFP and all three structural variability metrics were significantly correlated with the S-A axis. **e-g.** Age-related developmental trajectories of whole-brain mean coupling strength.

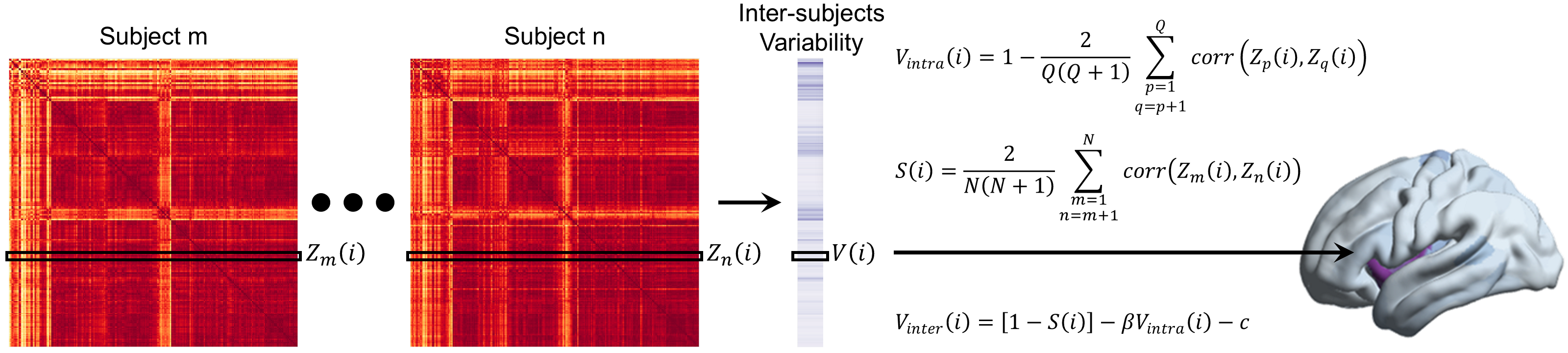

**Figure S10:** Flowchart of the vFC calculation process

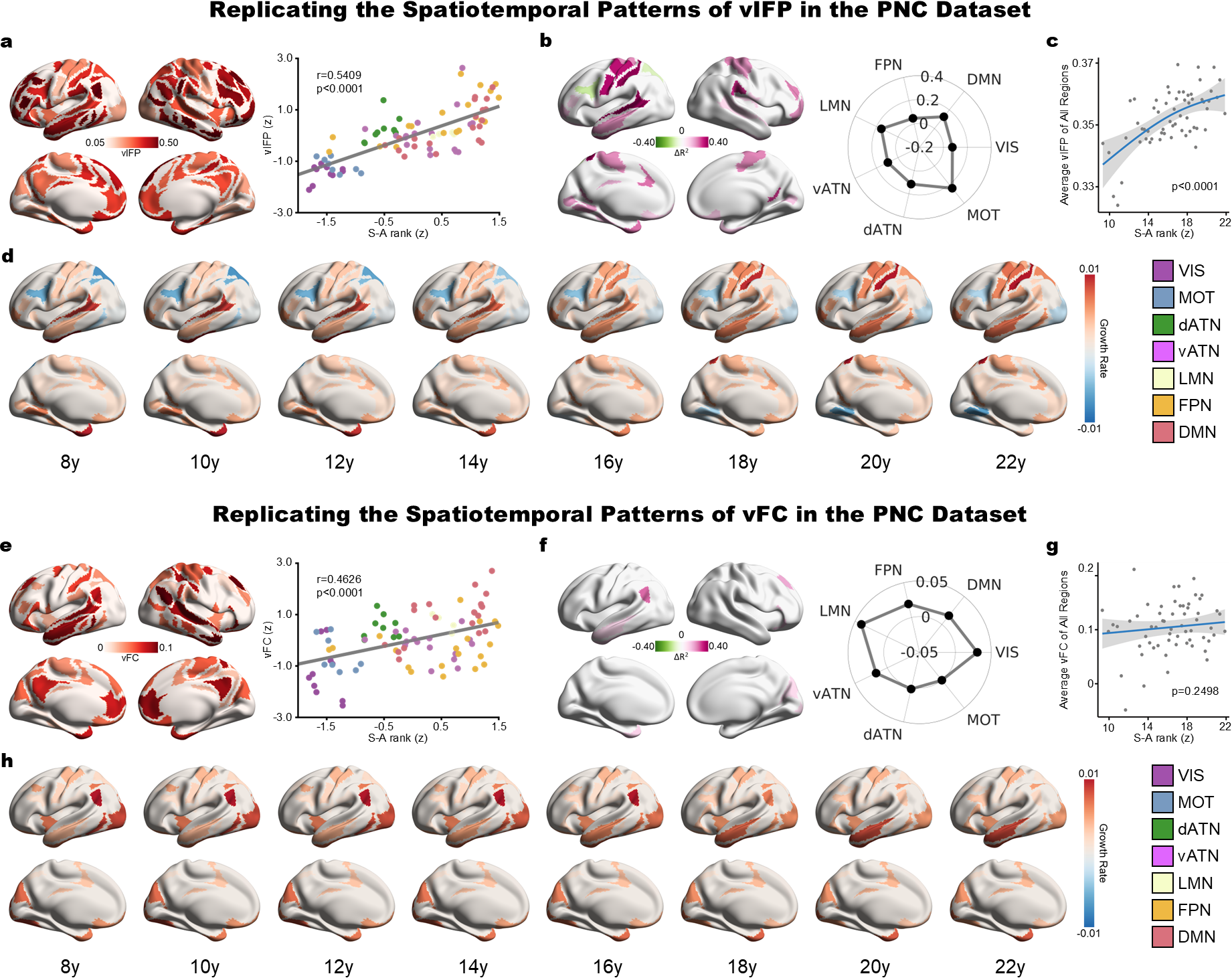

**Figure S11:** **Replication in the PNC dataset. a.** Spatial distribution of inter-individual variability in individualized functional parcellation (vIFP; n=610). Left: vIFP for 610 participants from the PNC dataset. Right: the correlation between S-A rank and vIFP, with point colors indicating functional networks. **b.** Regions with significant age effects on vIFP, with color indicating the magnitude and direction of the age effect. **c.** Developmental trajectory of average vIFP across the whole brain. **d.** The growth rate of vIFP was obtained as the first derivative (with respect to age) of the developmental trajectory fitted by a generalized additive model (GAM). **e.** Spatial distribution of inter-individual variability in functional connectivity (vFC; n=610). **f.** Regions with significant age effects on vFC, with color indicating the magnitude and direction of the age effect. **g.** Developmental trajectory of average vFC across the whole brain. h. Growth rate of vFC.
